# Cancer-specific loss of *TERT* activation sensitizes glioblastoma to DNA damage

**DOI:** 10.1101/2020.04.25.061606

**Authors:** Alexandra M. Amen, Christof Fellmann, Katarzyna M. Soczek, Shawn M. Ren, Rachel J. Lew, Gavin J. Knott, Jesslyn E. Park, Andrew M. McKinney, Andrew Mancini, Jennifer A. Doudna, Joseph F. Costello

**Author notes:** Correspondence should be addressed to J.A.D. and J.F.C. These authors contributed equally (Alexandra M. Amen, Christof Fellmann).

## Abstract

Most glioblastomas (GBMs) achieve cellular immortality by acquiring a mutation in the telomerase reverse transcriptase (*TERT*) promoter. *TERT* promoter mutations create a binding site for a GA binding protein (GABP) transcription factor complex, whose assembly at the promoter is associated with *TERT* reactivation and telomere maintenance. Here, we demonstrate increased binding of a specific GABPB1L isoform-containing complex to the mutant *TERT* promoter. Furthermore, we find that *TERT* promoter mutant GBM cells, unlike wild-type cells, exhibit a critical near-term dependence on GABPB1L for proliferation, both in cell culture and post-tumor establishment *in vivo*. Upregulation of the protein paralogue GABPB2, which is normally expressed at very low levels, can rescue this dependence. More importantly, when combined with frontline temozolomide (TMZ) chemotherapy, inducible GABPB1L knockdown and the associated *TERT* reduction led to an impaired DNA damage response that resulted in profoundly reduced growth of intracranial GBM tumors. Together, these findings provide new insights into the mechanism of cancer-specific *TERT* regulation, uncover rapid effects of GABPB1L-mediated *TERT* suppression in GBM maintenance, and establish GABPB1L inhibition in combination with chemotherapy as a novel therapeutic strategy for *TERT* promoter mutant GBM.

## Introduction

Primary glioblastoma (GBM) is the most common and lethal form of malignant brain cancer in adults. Current treatment strategies are limited, with GBM progression leading to death within two years of diagnosis in 90% of cases^1–3^. In GBM, as well as the vast majority of other cancers, transcriptional activation of the telomerase reverse transcriptase (*TERT*) gene, which is normally silenced in somatic cells, is a key step in tumorigenesis^4, 5^. *TERT* encodes the catalytic subunit of telomerase, and its reactivation in cancer is thought to contribute to cell survival and immortalization^5–8^. TERT ablation thus has the potential to directly affect both short- and long-term cell viability, through telomere length-dependent and independent pathways^9–15^. Prior research has demonstrated that inhibition of TERT expression enhances sensitivity of cells to DNA damage by radiation and chemotherapy, suggestive of a possible combination therapy for cancer treatment^16, 17^. However, telomerase inhibition is toxic to normal telomerase-dependent stem and germline cells, which has led to the failure of such approaches clinically^18–20^. Therefore, understanding genetic contributors to aberrant TERT expression, as well as consequences of cancer-specific TERT ablation, are critical to develop novel therapeutic avenues for GBM.

A major mechanism by which *TERT* is reactivated in cancer involves the acquisition of somatic mutations in its promoter, which comprise the most common non-coding mutations in cancer, and the third most common mutations overall^21–26^. In particular, ∼80% of primary GBMs contain one of two common single-nucleotide mutations that are associated with re-expression of *TERT* mRNA, referred to as G228A and G250A^22, 24, 27^. Both G-to-A transitions generate an identical 11 base pair sequence (plus strand: CCCGGAAGGGG) that creates a binding site for GA binding protein A (GABPA), an ETS-family transcription factor^28, 29^. Interestingly, these *de novo* ETS binding sites occur within three (G228A) or five (G250A) complete helical turns of two overlapping native ETS binding sites (ETS 195, ETS 200) in the *TERT* promoter (*TERT*p), one or the other of which are also required for *TERT* reactivation^28^. GABP transcription factors are obligate multimers that consist of the DNA-binding GABPA subunit (GeneID: 2551) and a transactivating GABPB subunit^30^. Humans have two paralogues encoding different beta subunits, GABPB1 (GeneID: 2553) and GABPB2 (GeneID: 126626). Reduced function of the long protein isoform of GABPB1 (GABPB1L) via indel mutations has previously been linked to downregulation of *TERT* mRNA and long-term telomere attrition^9^. This could provide a cancer-specific way to target TERT, particularly given that GABPB1L is dispensable for normal murine development, while GABPA and total GABPB1 are not^31, 32^. However, the extended time period required to induce cell death via progressive telomere shortening limits the therapeutic potential of this approach for high-grade GBM^33^. To further our understanding of the GABPB1L-TERT axis and its possible therapeutic benefit, here we examined the specificity and binding affinity of GABPB1L for the mutant *TERT* promoter, as well as functional effects of GABPB1L loss in a near-term, clinically relevant timeframe. Notably, we observed a dramatic synergistic effect between GABPB1L reduction and standard-of-care temozolomide chemotherapy, mediated through TERT downregulation and a resulting attenuation of the DNA damage response (DDR) in *TERT*p mutant cells. These rapid effects of targeting the cancer-specific GABPB1L-TERT axis in combination with frontline chemotherapy in established intracranial GBM xenografts led to substantially increased survival, promising exciting new avenues for *TERT*p-mutant GBM therapy.

## Results

### GABPB1L-containing complexes bind and regulate the mutant TERT promoter

*GABPB1* encodes two main transcript variants, a short isoform (GABPB1S) and a long isoform (GABPB1L). GABPB1S functions as a heterodimer with GABPA (GABPA_1_B_1_), while GABPB1L forms a heterotetramer (GABPA_2_B_2_) due to its unique terminal exon that contains a leucine zipper-like domain^30, 34^ (Fig. S1A). Previous work demonstrated that *TERT*p mutations create a binding site for the GABPA-B transcription factor complex, and linked reactivation of *TERT* in this context to GABP complexes containing GABPB1L^9, 28^. Indeed, the fact that two GABP binding sites distanced at complete helical turns from one another are required to enable *TERT* reactivation is suggestive of the recruitment of a GABPB1L-containing heterotetramer^9, 28, 35^ (Fig. S1A). However, GABPB1 isoform-specificity in mutant *TERT* promoter regulation has not been examined. In order to test this, we made use of a doxycycline-inducible, microRNA-embedded short hairpin RNA (shRNA) system with a green fluorescent protein (GFP) marker reporting shRNA induction^36–38^. We used a machine learning algorithm^38^ to design shRNAs targeting either GABPB1L or GABPB1S specifically (Fig. S1B). Doxycycline-mediated expression of GABPB1L or GABPB1S shRNAs led to reduction of the respective mRNA and protein levels in *TERT*p mutant (U-251) and *TERT*p wild-type (LN-18) GBM cells (Fig. 1A and Fig. S1C,D). Interestingly, GABPB1L – but not GABPB1S – knockdown led to reduced *TERT* mRNA and telomerase activity, exclusively in *TERT*p mutant cells (Fig. 1B,C and Fig. S1E), suggestive of *TERT*p mutant regulation of *TERT* expression through GABPB1L-containing complexes specifically. Of note, residual *TERT* expression and telomerase activity are observed in *TERT*p mutant cells following GABPB1L knockdown, when compared to a cancer cell line that is negative for *TERT* expression (U2OS) (Fig. 1B,C and Fig. S1E). This may be due to incomplete shRNA-mediated knockdown (Fig. 1A and Fig. S1C), or to regulation of *TERT* by other factors in conjunction with, or in the absence of, GABPB1L^39–41^.

**Fig. 1.**
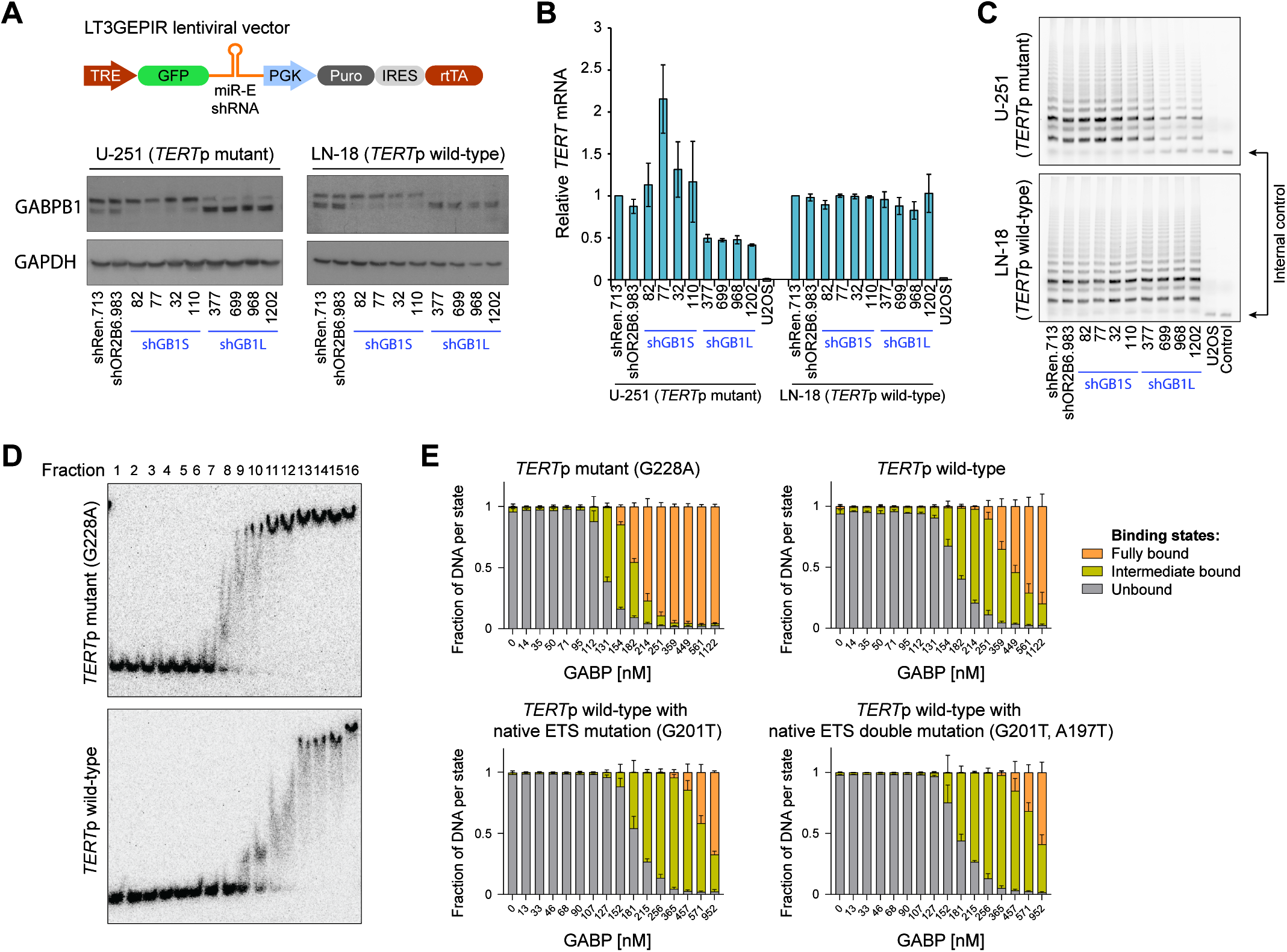
GABPB1L-containing complexes bind and regulate the mutant *TERT* promoter. **A**, Representative immunoblots of GABPB1 in U-251 and LN-18 cells expressing doxycycline-induced shRNAs targeting GABPB1S (shGB1S.82, shGB1S.77, shGB1S.32, shGB1S.110) and GABPB1L (shGB1L.377, shGB1L.699, shGB1L.968, shGB1L.1202) compared to negative control (olfactory receptor OR2B6, shOR2B6.910) and non-targeting (renilla luciferase, shRen.713) shRNAs. Cells were incubated with doxycycline for 6 days prior to harvest. Lower bands represent GABPB1S, upper bands represent GABPB1L. **B**, *TERT* mRNA expression measured via qRT-PCR in cell lines from (**A**), compared to a control cell line (U2OS) lacking *TERT* expression. **C**, Representative gels of telomerase activity measured via TRAP assay in cell lines from (**A**), compared to a control cell line (U2OS) lacking *TERT* expression. Control condition reflects no lysate. **D**,**E**, Representative gels (**D**) and quantification (**E**) of EMSAs comparing binding affinity of GABPA-B1L heterodimers to the mutant (G228A) *TERT* promoter (3 ETS binding sites), wild-type *TERT* promoter (2 native ETS binding sites), and control wild-type *TERT* promoter sequences lacking native ETS binding sites (G201T: native ETS single mutant; G201T, A197T: native ETS double mutant).

Though binding of GABP to the *TERT* promoter has previously been demonstrated^28^, possible GABPA_2_B1L_2_ heterotetramer formation at the mutant promoter has not directly been examined. To determine the nature of the GABPA-B1L interaction with the *TERT* promoter, we titrated purified heterodimer (Fig. S1F,G) against a fixed concentration of radiolabeled wild-type *TERT*p DNA, mutant *TERT*p DNA (G228A), and control probes with single (G201T) or double (A197T, G201T) mutations in the native ETS sites contained within the wild type promoter. We observed that GABPB1L-containing complexes bind with increased affinity to the mutant *TERT*p. However, calculating Kd values that accurately reflect GABP binding was challenging due to the observation of multiple intermediate states, which may be reflective of heterodimer binding, as has been previously suggested^42, 43^ (Fig. 1D and Fig. S1H). Thus, we segregated observed states into unbound, intermediate bound, and fully bound states. The comparison of signal ratios in each state for each target DNA tested across multiple GABP concentrations showed that the *TERT*p mutation (G228A) leads to increased formation of the fully bound state, which we attribute to a likely tetrameric complex^42, 43^ (Fig. 1D,E). We saw that GABP has the ability to bind all *TERT*p sequences tested in this study. However, binding to wild-type and native ETS site mutants (G201T or A197T, G201T) showed an increased number of intermediate states when compared to the G228A bearing promoters, and a decrease in the fully bound state (Fig. 1D,E and Fig. S1H). Together these results demonstrate that *TERT*p mutations result in increased binding affinity for probable GABPB1L-containing heterotetramer complexes, which help drive TERT reactivation.

### GABPB1L knockout impairs near-term growth of TERT promoter mutant GBM cells

For GBM therapy, short-term effects of *TERT* reduction, rather than long-term consequences of canonical telomere attrition, would increase the clinical relevance of targeting the cancer-specific GABP-TERT axis^33^. Given that GABPB1L regulates *TERT* in the context of *TERT*p mutations, our data raise the question of possible early functional consequences of GABPB1L inhibition on *TERT*p-mutant cell growth and viability. CRISPR-Cas methods enable the precise dissection of genetic dependencies through the generation and analysis of specific heterozygous and homozygous genetic deletions in isogenic cell lines^44^. Therefore, to assess near-term consequences of complete GABPB1L deletion, we used CRISPR-Cas9 and single-guide RNAs (sgRNAs) targeting the intron and 3’UTR flanking the unique exon 9 of *GABPB1L* (Fig. 2A, Fig. S2A,B, and Table S1). We compared a set of three *TERT*p mutant GBM cell lines (U-251, LN-229, and T98G) and four *TERT*p WT cell lines (HEK293T, HAP1, NHA-S2, and LN-18) by generating monoclonal isogenic derivatives. The *TERT*p WT lines include an immortalized human astrocyte line (NHA-S2) as well as a GBM line (LN-18). Sequencing of the *TERT* promoter confirmed *TERT*p mutation status (Fig. S2 C,D).

**Fig. 2.**
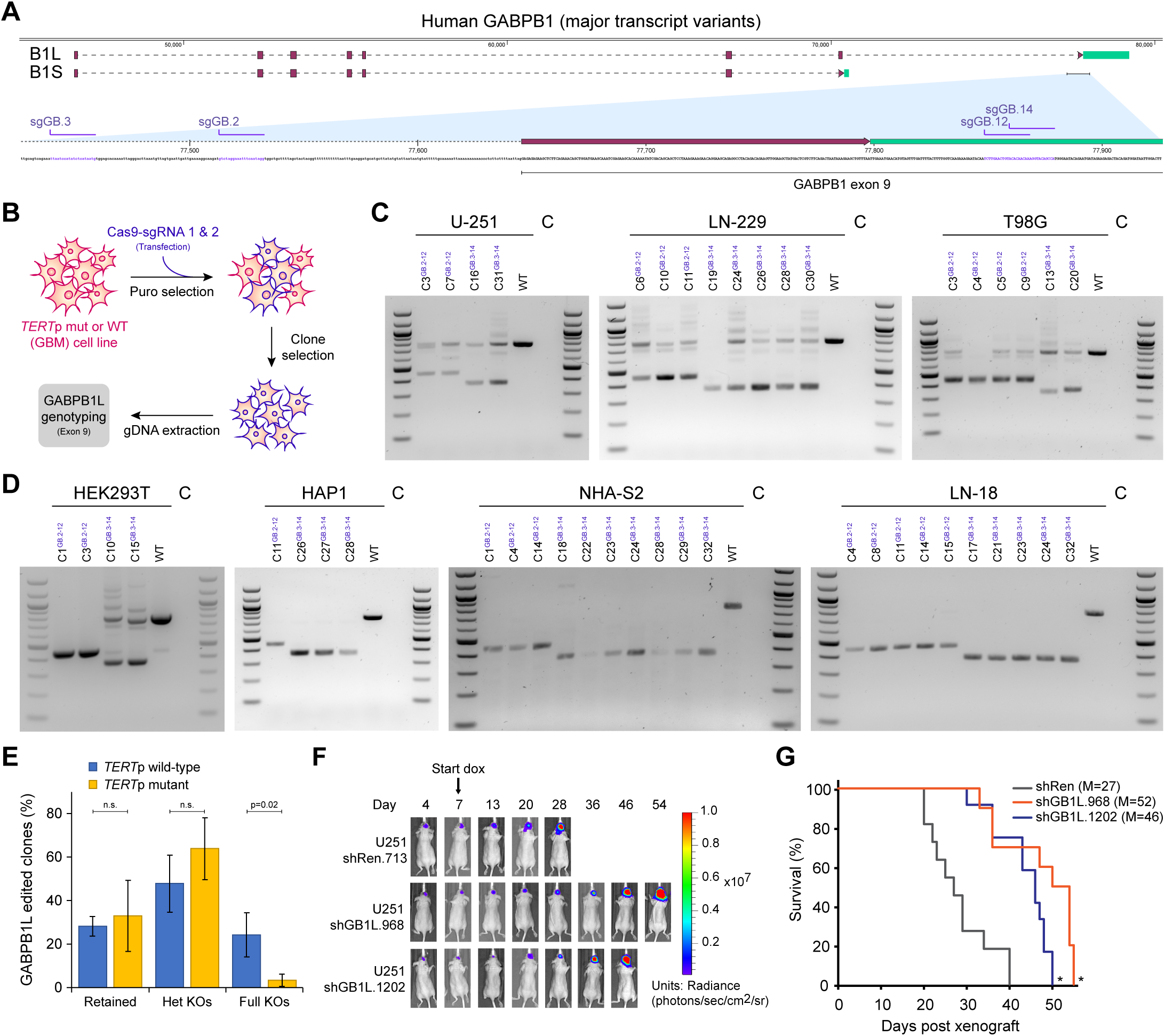
Inhibition of GABPB1L rapidly prevents growth of *TERT* promoter mutant cells. **A**, *GABPB1L* exon 9 excision strategy. Diagram of the major human *GABPB1* transcript variants, encoding either a short (*GABPB1S*) or long (*GABPB1L*) isoform. Red rectangles represent the coding sequence (CDS), green rectangles the 3’ untranslated region (3’UTR). For clarity, the first exon containing the 5’ untranslated region (5’UTR) is not shown. *GABPB1L* sgRNA targeting sites are highlighted in purple. **B**, Editing strategy for scarless knockout of the long isoform of GABPB1 (GABPB1L). In each case, a pair of sgRNAs were used targeting *GABPB1L* intron number 8 and the 3’ UTR of *GABPB1L*, respectively. **C**,**D**, Genotyping analysis of *GABPB1L* edited monoclonal *TERT* promoter mutant U-251, LN-229 and T98G GBM cell lines (**C**) and *TERT* promoter WT HEK293T, HAP1, NHA-S2 and LN-18 cell lines (**D**). Superscripts indicate the sgRNA pair used for editing. WT: parental wild-type cell line. C: PCR no template control. **E**, Comparison of *GABPB1L* editing outcomes between *TERT*p wild-type (blue) and *TERT*p mutant (yellow) lines. Bars represent the percentage of each editing outcome averaged over 3 *TERT*p mutant and 4 *TERT*p wild-type lines. Retained refers to clones where the *GABPB1L* exon 9 was overall retained, even though they may contain indels at either or both of the sgRNA target sites flanking exon 9. p: p-value (unpaired, two-tailed student’s t-test). n.s.: not significant (alpha level = 0.05). **F,G,** Representative bioluminescence images (**F**) and Kaplan-Meier survival curve (**G**) of mice injected with U-251 cells expressing a control shRNA (shRen.713) or GABPB1L-targeting shRNAs (shGB1L.968, shGB1L.1202). Mice were fed doxycycline chow to induce shRNA expression starting at day 7 post-injection. n=10-12 mice per condition. *: p-value<0.01 compared to shRen.713 (Kaplan-Meyer log-rank test). M: median survival.

To identify ideal guide RNAs, all possible combinations of three intronic and three 3’UTR targeting guides were compared in HEK293T and U-251 cells to assess editing efficiency (Fig. 2B and Fig. S2E). Next, the two most efficient pairs of sgRNAs were used to edit all seven cell lines, and 169 monoclonal cell line derivatives were established, genotyped, and subjected to further analysis (Fig. 2C,D, Fig. S3A-H, and Fig. S4A-C). Strikingly, we observed that while full homozygous knockout of GABPB1L occurred at an appreciable rate across *TERT*p WT cell lines (13-36%), we were able to generate only one full GABPB1L knockout line, LN229-C19, among the 76 *TERT*p mutant clones that were screened (Fig. 2E and Fig. S3A-H). Importantly, the discrepancy between *TERT*p WT and mutant cell lines did not appear to result from differences in overall editing efficacy between cell lines, as equal numbers of heterozygous knockouts were observed (Fig. 2E and Fig. S3H). Prior studies from our lab demonstrated that impaired GABPB1L protein function through indel-based editing led to long-term telomere attrition over a more than 90-day timeframe^9^. However, the present data suggest that full knockout of GABPB1L causes *TERT*p mutant cells to become either slow-growing or nonviable over the ∼30 day editing and selection process, a timeframe that is not consistant with gradual telomere shortening^9, 15, 45^.

### GABPB1L knockdown slows growth of established TERTp mutant tumors

To more directly assess the therapeutic potential of GABPB1L reduction, we next examined its effect on growth of established GBM tumors *in vivo*, using the above-described doxycycline-inducible shRNA system (Fig. 1A). We first tested the system for *in vivo* efficacy using a *bona fide* essential gene, Replication Protein A1 (RPA1, GeneID: 6117), along with controls. Addition of doxycycline to U-251 cells stably transduced with an inducible vector encoding RPA1-targeting shRNAs resulted in reduced *RPA1* mRNA *in vitro* (Fig. S5A) and prolonged animal survival *in vivo* (Fig. S5B,C). Because the doxycycline-inducible vectors encode an shRNA within the 3’ UTR of a GFP construct (Fig. 1A), the percentage of GFP-positive cells was used as a proxy readout for shRNA expression. *Post mortem* tumor analysis showed that *in vivo* expression of the RPA1-targeting shRNA construct was reduced compared to controls (Fig. S5D), consistent with a survival advantage for loss of RPA1 knockdown^37^.

We then made use of our GABPB1L-targeting shRNAs to determine the effect of GABPB1L knockdown on growth of existing GBM tumors. Two separate GABPB1L-targeting shRNAs tested in orthotopic intracranial xenografts of U-251 cells slowed tumor growth within 12 days of shRNA induction when induced post tumor implantation (Fig. 2F and Fig. S5E), and increased median survival by 70-90% (Fig. 2G). GABPB1L shRNA-mediated target knockdown and linked GFP expression were maintained *in vivo*, via analysis of *ex vivo* dissociated tumor cells *post mortem* (Fig. S5F,G). A third tested GABPB1L-targeting shRNA (shGB1L.699) did not impact survival (Fig. S5H). Correspondingly, *post mortem* analysis demonstrated minimal GABPB1L protein reduction in tumors with this shRNA, and showed that only ∼30% of tumor cells retained GFP expression (Fig. S5F,G), indicating that sustained GABPB1L knockdown is necessary to slow tumor growth. Together, these data demonstrate that shRNA-mediated GABPB1L suppression in established tumors significantly prolongs animal survival, highlighting its therapeutic value.

### GABPB2 promotes resistance to GABPB1L loss in TERTp mutant GBM cells

Given the potential therapeutic benefit of GABPB1L inhibition in *TERT*p mutant GBM, we chose to analyze heterozygous knockout clones generated from *TERT*p mutant lines compared to full and heterozygous knockouts from *TERT*p WT lines, in order to better understand the molecular impact of GABPB1L loss (Fig. 2C-E). As expected, full or heterozygous GABPB1L knockout led to reduction or loss of *GABPB1L* mRNA and protein in all cell lines (Fig. S6A,B). We also observed unchanged or upregulated total *GABPB1* mRNA (all transcript variants), and substantially upregulated *GABPB1S* mRNA and protein across all cell lines (Fig. S6A-F), supporting the possibility that all *GABPB1* transcripts from a *GABPB1L* knockout locus are now *GABPB1S*. Notably, heterozygous GABPB1L knockout was associated with a decrease in *TERT* mRNA expression and reduced telomerase activity in *TERT*p mutant cells only (Fig. S6G-J), confirming regulation of *TERT* by GABPB1L in a *TERT*p mutant-specific manner.

The single *TERT*p mutant GABPB1L full knockout cell line (LN229-C19) that we obtained did not exhibit reduction of either *TERT* mRNA or telomerase activity (Fig. S5G,I,J), suggesting a compensatory mechanism to regulate *TERT* independent of GABPB1L. *GABPB1* and *GABPB2* are distinct genes with high sequence conservation that perform similar but separate functions, though GABPB2 is thought to function exclusively as a heterotetramer with GABPA^30, 34^, mediated by the presence of a leucine zipper-like domain (Fig. 3A, Fig. S7A). Since GABPB2 is expressed at lower levels than GABPB1 in GBM based on data from the TCGA Research Network (https://www.cancer.gov/tcga) (Fig. S7B), we probed for possible GABPB2 upregulation in our GABPB1L knockout clones. While deletion of GABPB1L did not affect *GABPB2* mRNA in any other line, the LN229-C19 clone showed a 4.5-fold elevation in *GABPB2* transcript levels (Fig. 3B and Fig. S7C). In support of a potential compensatory mechanism via GABPB2 upregulation, stable GABPB2 overexpression in U-251 cells rescued cell dropout following CRISPR editing of the *GABPB1* locus (Fig. 3C). In addition, small interfering RNA (siRNA)-mediated inhibition of *GABPB2* showed that *TERT* mRNA levels are responsive to GABPB2 knockdown in LN229-C19, but not in wild-type LN-229 cells or heterozygous knockout clones (Fig. 3D). These data suggest that GABPB2 upregulation can compensate functionally for *GABPB1L* deletion in *TERT*p mutant cells, and indicate that *TERT*p mutant GBM can tolerate total loss of GABPB1L in the context of alternative mechanisms rescuing TERT expression. This finding reveals a possible resistance mechanism for GABPB1L-targeting therapies.

**Fig. 3.**
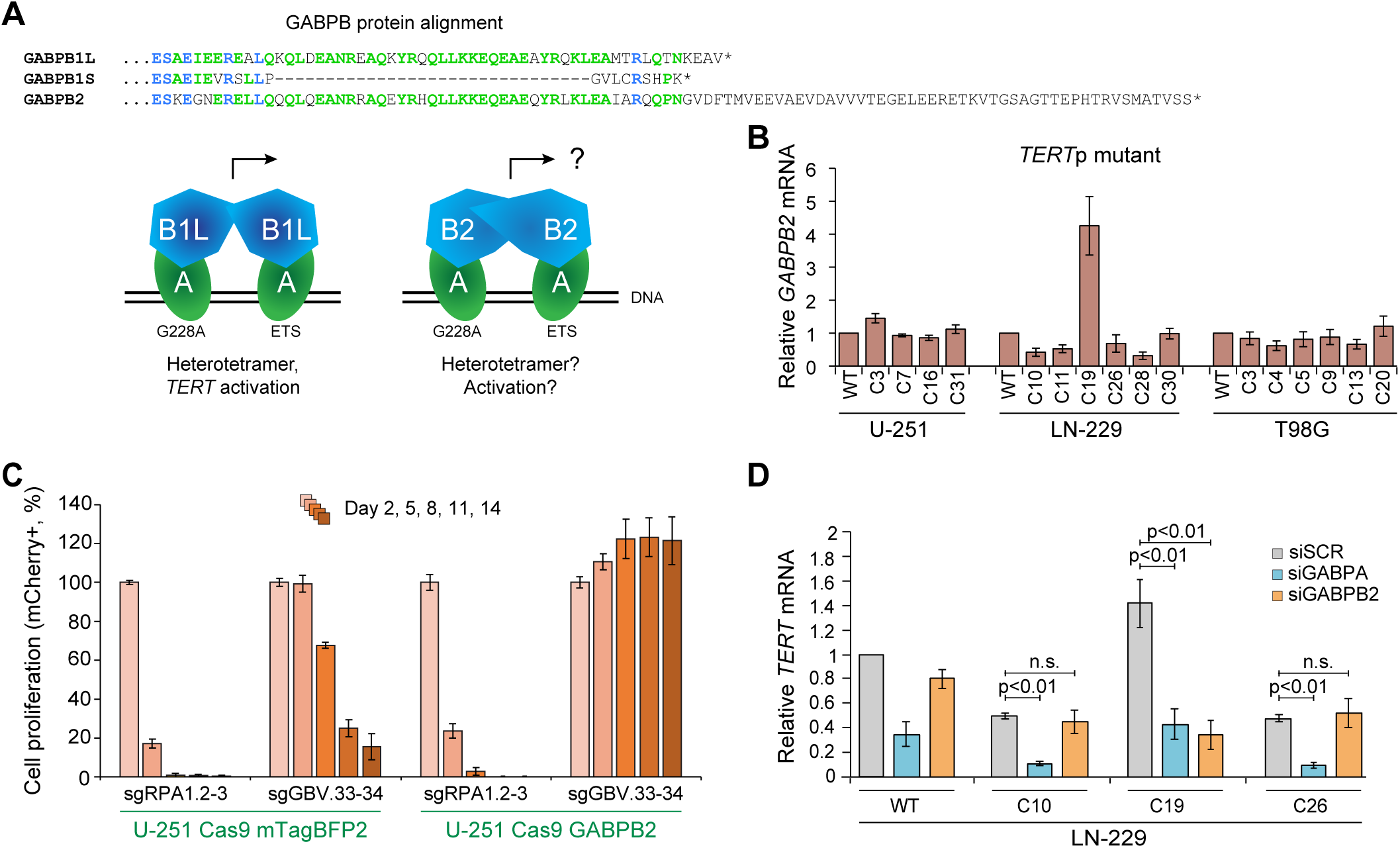
GABPB2 upregulation can compensate for loss of GABPB1L. **A**, Similarity between GABP proteins. Partial protein alignment (MUSCLE) of GABPB1L, GABPB1S, and GABPB2. GABPB1L is the long isoform, and GABPB1S the short isoform, of the *GABPB1* gene. *GABPB2* is a distinct gene. Amino acids matching across all three proteins are highlighted in blue, those matching across two proteins are highlighted in green. GABPA (alpha) subunits of the GABP complex bind to both the native ETS site (ETS) and the mutation-derived ETS sites (G228A or G250A) at the TERT promoter locus. The GABPB1 short (B1S) or long (B1L) isoform subunits bind to the alpha subunits to form either heterodimers (GABPA_1_B1S_1_) or heterotetramers (GABPBA_2_B1L_2_). Heterotetramer formation is presumably mediated through the leucine zipper-like domain of GABPB1L, which is also present in GABPB2. **B,** *GABPB2* mRNA expression measured via qRT-PCR in GABPB1L knockout clones, plotted relative to control (wild-type) cells, in *TERT*p mutant cell lines. Data represent mean +/- SEM. **C,** Competitive proliferation assay in U-251 cells using pairs of sgRNAs targeting a positive control locus (sgRPA1.2-3) and total GABPB1 (targeting exon 3, sgGABPV.33-34), in the presence of a lentiviral vector expressing either GABPB2 or mTagBFP2 (control). Data represent mean +/- standard deviation of triplicates. **D**, *TERT* mRNA expression measured via qRT-PCR following siRNA-mediated knockdown of GABPA or GABPB2 in LN-229 wild-type, FKO, or heterozygous KO cells. p: p-value (unpaired, two-tailed student’s t-test). n.s.: not significant (alpha level = 0.05). Data represent mean +/- SEM.

### Increased response of GABPB1L-reduced GBM tumors to chemotherapy

Currently, most patients diagnosed with primary GBM are treated with a regimen that includes temozolomide (TMZ) chemotherapy, a DNA alkylating chemotherapeutic agent that is most effective at eliminating cells that lack O-6-methylguanine-DNA methyltransferase (MGMT) through generation of single- and double-strand DNA breaks^46, 47^. Prior studies have indicated that reduction of *TERT* in *TERT*-expressing cells may sensitize cells to DNA damage from ionizing radiation and chemotherapy^16, 17^. Reduction of *TERT* mRNA by RNAi has been shown to prevent the repair of DNA breaks *in vitro* by inhibiting central components of the DNA damage response (DDR), such as histone H2AX phosphorylation (yH2AX), through an incompletely understood mechanism^16^. Loss of yH2AX-mediated DNA damage signaling increases DNA damage-related senescence or cell death, possibly through a reduction of normal cell cycle arrest^16, 48, 49^. Therefore, we wondered whether GABPB1L-mediated reduction of *TERT* mRNA in *TERT*p mutant GBM might promote TMZ response, enabling a possible path to GBM cell-specific chemotherapy sensitization (Fig. 4A).

**Fig. 4.**
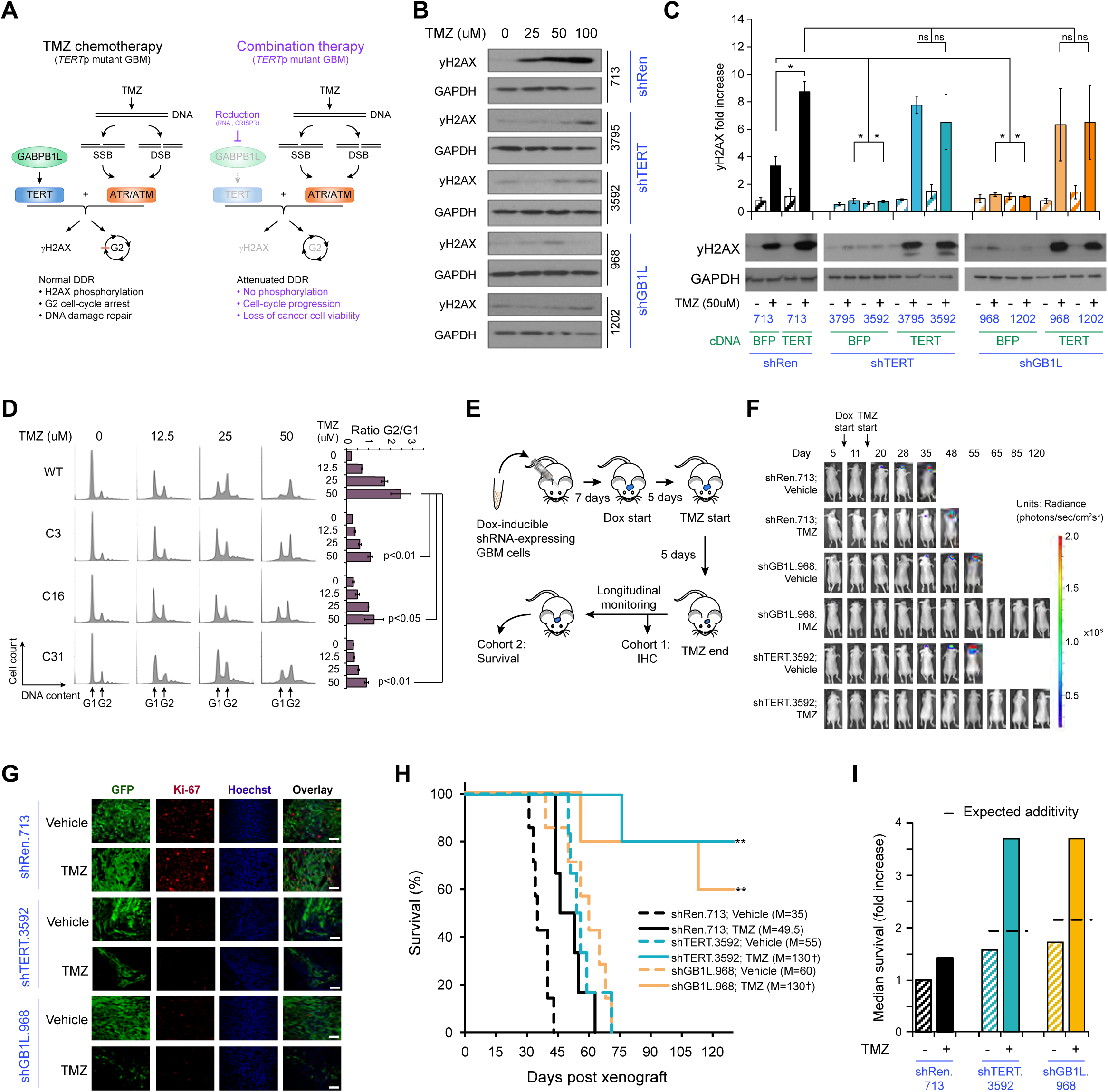
Reduction of GABPB1L potentiates anti-tumor response of *TERT*p mutant GBM to chemotherapy. **A,** Schematic for possible sensitization of *TERT* promoter mutant GBM cells to TMZ through GABPB1L inhibition. Inhibiting GABPB1L leads to reduced *TERT* expression, resulting in a blunted DNA damage response and lack of normal cell cycle arrest, ultimately reducing cancer cell viability following DNA damage. SSB: single-strand break. DSB: double strand break. DDR: DNA-damage response. **B**, Representative immunoblots of yH2AX in U-251 cells expressing doxycycline-induced shRNAs targeting TERT (shTERT.3795, shTERT.3592) or GABPB1L (shGB1L.968, shGB1L.1202) compared to a non-targeting (renilla luciferase, shRen.713) shRNA. **C**, Representative immunoblots (bottom) and quantification (top, from triplicate blots) of yH2AX in U-251 cells engineered to stably express either BFP (control) or TERT (rescue) in the presence of shRNAs targeting TERT (shTERT.3795, shTERT.3592) or GABPB1L (shGB1L.968, shGB1L.1202) compared to a non-targeting shRNA (renilla luciferase, shRen.713). **B**,**C**, Cells were incubated with doxycycline for 6 days prior to harvest, and treated with specified dose(s) of TMZ 20 hours prior to harvest. **D**, Representative images (left) and G2/G1 ratio quantification (right) from FACS-based cell cycle analysis of U-251 wild-type and GABPB1L heterozygous knockout cells treated with a dose titration of TMZ 72 hours prior to harvest. p: p-value (unpaired two-tailed student’s t-test). **E**, Schematic of *in vivo* temozolomide (TMZ) treatment. Mice were xenografted with U-251 cells expressing doxycycline-inducible shRNAs, and placed on doxycycline chow 7 days post orthotopic xenograft. Mice were then treated with TMZ or a vehicle control by oral gavage starting 12 days post-xenograft for 5 days. IHC: immunohistochemistry. **F,** Bioluminescence imaging of mice injected with U-251 cells engineered to express shRNAs targeting TERT (shTERT.3592), GABPB1L (shGB1L.968), or a non-targeting control shRNA (shRen.713). Mice were placed on doxycline chow and p.d. dosed with either TMZ or a vehicle control as described in (**E**). **G**, Representative images of immunofluorescence staining in tumors from mice described in (**E**), cohort 1. Mice were euthanized 30 days post-orthotopic xenograft for analysis. Doxycycline-induced tumor cells are GFP positive. N=5 images per mouse, 3 mice per condition. Scale bar = 50uM. **H**, Kaplan-Meier survival curves for mice described in (**E**), cohort 2. For statistics, survival of mice with no tumor burden was set at the experimental endpoint (130 days). N = 5-7 mice per condition. **: p-value<0.01, relative to all vehicle conditions (Kaplan-Meier log-rank test). M: median survival. **I**, Comparison of the relative fold-increase in median survival based on overall survival in (**H**). Dashed line represents a simple addition of effects of TMZ plus GABPB1L or TERT knockdown. Survival of mice xenografted with wild-type cells and treated with vehicle control was used for normalization.

U-251 cells expressing doxycycline-inducible shRNAs targeting either TERT or GABPB1L were treated with a dose titration of TMZ, and the DDR was assessed through analysis of yH2AX levels. Compared to control, shRNA-mediated TERT knockdown significantly reduced upregulation of yH2AX 20 hours after treatment with TMZ (Fig. 4B and Fig. S8A,B). Strikingly, GABPB1L knockdown led to an equally blunted yH2AX DDR following TMZ treatment (Fig. 4B and Fig. S8B). Similarly, TMZ-mediated yH2AX upregulation was reduced in both U-251 and LN-229 clones with heterozygous GABPB1L knockout, compared to wild-type cells (Fig. S8C,D). However, the full GABPB1L knockout clone LN229-C19 with GABPB2 upregulation and maintained TERT expression exhibited TMZ-induced yH2AX induction that mirrored wild-type, consistent with the impairment of the DDR response being mediated by TERT downregulation (Fig. S8D).

To directly assess whether GABPB1L knockdown leads to reduced yH2AX upregulation through reduced TERT, we generated U-251 cells that stably express either BFP (control) or TERT (rescue) constructs (Fig. S8E). While the BFP construct showed no effects, overexpression of TERT completely rescued loss of yH2AX upregulation following shRNA-mediated knockdown of both TERT and GABPB1L (Fig. 4C). In the control shRNA condition (shRen.713), TERT overexpression significantly increased yH2AX upregulation following TMZ treatment when compared to wild-type TERT levels, providing further evidence for regulation of the DDR by TERT (Fig. 4C and Fig. S8E).

Ablated DDR signaling can lead to a blunting of the G2 cell cycle arrest that is commonly observed following treatment with DNA damaging agents such as TMZ, causing an increased loss of viability as cells continue to divide^48, 50, 51^. In concordance, here we also observe reduced G2 arrest in both U-251 and LN-229 GABPB1L heterozygous knockout clones treated with TMZ, as compared to both wild-type cells and the LN-229 full GABPB1L knockout clone C19 (Fig. 4D, Fig. S8F,G). Lack of MGMT expression was confirmed in all cell lines and corresponding clones (Fig. S8H).

Both partial and full loss of H2AX, or inhibition of its phosphorylation, have been shown to significantly impact cell survival following exposure to DNA damage^52–54^. Similarly, inhibition of G2 arrest following TMZ administration is associated with decreased cell viability and growth, resulting in TMZ sensitization^50^. Therefore, we hypothesized that *TERT*p mutant GBM cells with reduced GABPB1L would be more sensitive to TMZ chemotherapy. We first assessed the therapeutic implications of this finding *in vivo* through administration of a TMZ regimen following orthotopic intracranial xenograft of either U-251 or LN-229 wild-type and GABPB1L knockout cells. Initial analysis – in the absence of TMZ therapy – of GABPB1L heterozygous knockout cell tumors *in vivo* demonstrated that GABPB1L reduction slowed growth of both U-251 and LN-229 tumors and increased animal survival (Fig. S8I-L), while the full knockout LN229-C19 clone with maintained *TERT* expression formed tumors at a similar rate to WT cells. Strikingly, *in vivo* administration of chemotherapy in combination with GABPB1L heterozygous status radically ablated tumor growth of U-251 clones (Fig. S9A-D). A similar potentiation of treatment efficacy was also observed, though to a lesser extent, in GABPB1L-reduced LN-229 tumors, again with the expected exception of the full knockout LN229-C19 clone that maintained wild-type *TERT* mRNA levels (Fig. S9E-H).

Lastly, in order to assess the therapeutic potential of this combination therapy in a more clinically relevant manner, we performed orthotopic xenografts of U-251 cells engineered to express doxycycline-inducible shRNAs targeting either TERT (shTERT.3952) or GABPB1L (shGB1L.968), compared to a control. Following tumor engraftement, mice were placed on doxycycline and treated with a regimen of either TMZ or vehicle, then either sacrificed 30 days post-xenograft for immunohistochemistry analysis (Cohort 1), or followed to endpoint for survival analysis (Cohort 2) (Fig. 4E). We verified that doxycycline and TMZ do not interfere with each other to affect either tumor growth or shRNA induction *in vivo* (Fig. S9I,J). Interestingly, knockdown of TERT and GABPB1L both slowed growth of tumors and prolonged survival to a similar extent, which was accompanied by a significant reduction in the percentage of Ki-67 positive tumor cells in these conditions (Fig. 4F-H and Fig. S9K-M). More importantly, both TERT and GABPB1L reduction in established intracranial xenografts resulted in a similar potentiation of TMZ efficacy, such that tumor growth was either dramatically slowed or eliminated (Fig. 4H and Fig. S9K,L). The median animal survival was increased beyond a simple addition of effects (Fig. 4I), showing the signature of a synergistic combination therapy. Ki-67 status was not affected by TMZ treatment, as assessed 14 days following the TMZ regimen (Fig. 4G and Fig. S9M). Together, the dramatic anti-tumor effects of TMZ combined with GABPB1L reduction and the associated cancer-specific downregulation of *TERT* mRNA are of significant clinical interest, as they provide a path to increased TMZ response for *TERT*p mutant GBMs. Indeed, our data suggest that *MGMT*-silenced *TERT*p mutant GBM could be effectively targeted through a combination of GABPB1L reduction and TMZ treatment.

## Discussion

A major challenge in glioblastoma therapy is finding efficient methods to reduce growth of established tumors while leaving normal cells unaffected. Here, our data directly demonstrate increased binding of GABPB1L isoform-containing complexes to the mutant compared to wild-type *TERT* promoter. Additionally, we show that knockdown of GABPB1L results in near-term anti-growth effects on *TERT*p mutant cells both *in vitro* and *in vivo*, and that upregulation of GABPB2 can compensate for the loss of GABPB1L in this context. Mechanistically, the early appearance of these anti-growth effects hint towards non-canonical functions of TERT, rather than telomere attrition or immediate vulnerability of cells with short telomeres, though this warrants further investigation^9–11, 15, 45^. Additionally, these effects may be the direct result of TERT downregulation following GABPB1L reduction, a combined effect between reduced TERT and other factors associated with GABPB1L loss, or involve other differences in gene expression specific to *TERT*p mutant cells^55^. Regardless, our results provide the first evidence that even after tumor initiation, GABPB1L knockdown rapidly impairs growth of GBM tumors *in vivo*, which is of great therapeutic interest.

More strikingly, we also find that GABPB1L reduction combined with TMZ treatment dramatically potentiates anti-tumor effects *in vivo* through cancer-specific loss of TERT, emphasizing the clinical potential of this combination therapy. Specific targeting of TERT through the long isoform of GABPB1 (GABPB1L) is particularly promising given that both GABPA and total GABPB1 (the short and long isoform together) are required for normal murine development, while the GABPB1L isoform is dispensable^9, 31, 32^. Previous studies have observed that TERT reduction sensitizes cells to chemotherapy and radiation by inhibiting the normal DNA damage response (DDR), measured in part by a loss of yH2AX upregulation^16^. Here, our data show that reducing TERT through knockdown of GABPB1L prevents the upregulation of yH2AX and G2 cell-cycle arrest normally seen following exposure of *TERT*p mutant cells to standard-of-care TMZ chemotherapy. The mechanism of action of TMZ is largely through mismatch repair pathway-associated single strand breaks, but also results in DNA double strand breaks, and yH2AX signaling may occur in response to single-as well as canonically to double-strand breaks^46, 47, 56, 57^. Loss of yH2AX signaling leads to reduced cell viability following DNA damage induced by radiation and chemotherapy^48, 49, 54^. Our findings extend this by demonstrating that *TERT*p mutant GBM tumors with reduced *TERT* mRNA, either through knockdown of GABPB1L or by targeting TERT directly, show a dramatically increased response to TMZ *in vivo*.

Targeted cancer therapies have the potential to be highly impactful, but cancer cells commonly develop resistance to anti-tumor drugs^58^. Uncovering mechanisms of resistance often allows development of second-generation therapies that stunt tumor relapse. Our work points to sporadic GABPB2 upregulation as a possible resistance mechanism to loss of GABPB1L, providing preliminary evidence that GABPB2, which is typically expressed only at very low levels in GBM, could be targeted in conjunction with GABPB1L for increased treatment robustness. Multiple other transcription factors have also been linked to TERT regulation, both in normal and oncogenic contexts^59–62^. Such factors could play an additional role in basal expression of TERT from the mutant promoter, or compensate for loss of GABPB1L, and merit future study. While clinical methods to inhibit GABPB1L do not currently exist, and transcription factors such as the GABP complex are generally considered difficult to target, viable pathways might include approaches such as anti-sense oligo therapy, which has been successful at targeting transcription factors through regulation of splicing^63^, or development of small molecules that prevent GABPA_2_B_2_ heterotetramer formation. The possible clinical benefit of these approaches warrants further study. Together, our data suggest that GABPB1L inhibition combined with TMZ treatment can provide a tumor-specific path to improve disease outcomes for patients with *TERT*p mutant GBMs, and possibly the many other cancers harboring *TERT*p mutations.

## Methods

### Protein purification

GABPB1L and GABPA subunits were cloned into pET-Duet-1 vector with gibson assembly. The GABPB1L subunit was inserted after the first ribosome binding site and a plasmid encoded HisTag. HisTag and GABPB1L were separated by insertion of TEV cleavage sequence. GABPA was inserted after the second ribosome binding site without any affinity purification tags.

GABPB1L and GABPA were expressed in Rosetta DE3 cells. A single colony was used for an overnight starter culture in Luria-Bertani media (LB) with 100 µg/ml Ampicillin. Next, bacteria from starter culture was grown in Terrific Broth (TB) media with 100 µg/ml of Ampicillin at 37°C until OD_600_ 0.8-1. Cells were chilled on ice and induced with 0.5 mM IPTG and grown overnight at 16°C. Cells were harvested via centrifugation, resuspended in 50 mM HEPES pH 7 500 mM NaCl, 1 mM TCEP, 5% (v/v) glycerol and flash frozen. Cell pellets were thawed and 0.5 mM PMSF and cOmplete protease inhibitor (Roche) were added. Cells were lysed by sonication and lysates were cleared by centrifugation. GABPA-B1L heterodimers were purified first through Ni-NTA Superflow resin (Qiagen). Column was equilibrated in 50 mM HEPES pH 7, 500 mM NaCl, 1mM TCEP, 5% glycerol, 10mM imidazole. Cleared lysate was incubated with resin for 30 min on a rocker at 4°C. Resin was then washed with the above equilibration buffer and protein was eluted with 50 mM HEPES pH 7, 500 mM NaCl, 1 mM TCEP, 5% (v/v) glycerol, 500 mM imidazole. Fractions containing both subunits were combined and dialysed overnight in Slide-A-Lyzer (ThermoFisher) with addition of TEV protease in 50 mM HEPES pH 7.5, 150 mM NaCl, 1 mM TCEP, 5% (v/v) glycerol. Heterodimers were then purified through Ni HiTrap HP (GE) and Heparin HiTrap column (GE) or Q HiTrap HP using dialysis buffer as buffer A and 50 mM HEPES pH 7.5 1 M KCl, 1 mM TCEP, 5% (v/v) glycerol as eluting buffer B. Fractions containing heterodimers were combined, concentrated and loaded on Superdex 200 Increase 10/300 equilibrated with 20 mM HEPES pH 7.5, 200 mM KCl, 1 mM TCEP, 5% (v/v) glycerol. Purified protein was concentrated and flash frozen for storage at -80°C.

### Electrophoretic Mobility Shift Assay

S/ingle stranded 60bp oligonucleotides with wild type *TERT*p including native ETS sites (5’-AGGGCGGGGCCG**CGGAAAGGAA**GGGGAGGGGCTGGGAGGGCCCGGAGGGGGCTG GGCCGG - 3’), mutant *TERT*p with native and *de novo* ETS sites created by G228A mutation (5’- AGGGCGGGGCCG**CGGAAAGGAA**GGGGAGGGGCTGGGAGGGCC**CGGA<u>a</u>**GGG GCTGGGCCGG - 3’) and native ETS mutant sequences G201T (5’ – AGGGCGGGGCCG **CGGAAAGtAA**GGGGAGGGGCTGGGAGGGCCCGGAGGGGGCTGGGCCGG - 3’) and A197T, G201T (5’ – AGGGCGGGGCCG**CGGtAAGtAA**GGGGAGGGGCTGGGAGGGCCC GGAGGGGGCTGGGCCGG - 3’) were obtained from IDT and were Urea - PAGE purified, ethanol precipitated and resuspended in DEPC. One of the strands of each sequence was then labeled with P^32^ as follows: 10 pmols of single stranded DNA was incubated with 5 Units of T4 PNK (NEB) and 0.64µCi of P^32^- γ -ATP (Perkin-Elmer) for wild type and G228A mutant DNA and with 10 µCi for the G201T and A197T, G201T mutants in PNK buffer in total volume of 25µl for 30 min at 37°C and heat inactivated at 65°C for 20min. Reaction was diluted by addition of 25µl of DEPC. Excess of ATP was removed on desalt G25 (GE) column. Purified oligos were then further diluted by addition of 50µl of DEPC. Labeled oligos were annealed with corresponding unlabeled oligo in 20 mM Tris pH 7.5, 100 mM LiCl, 1 mM MgCl_2_. These double stranded substrates were used in binding experiments with GABPα

GABP heterodimers were thawed and diluted to stock concentration of 2 µM, which was further serially diluted to create additional 15 protein stocks in Superdex 200 buffer. DNA oligonucleotides were diluted to 1 nM stock concentration in annealing buffer. Reactions were performed by mixing 0.2 nM 60bp DNA with equal volumes of increasing concentrations of /β heterodimers (0 to ∼1000nM) with 5x reaction buffer (100 mM Tris pH7.5, 50 mM KCl, 25 mM MgCl_2_, 5 mM TCEP, 25% (v/v) glycerol in 15 µl final reaction volume. Reactions were incubated at room temperature for 1 hr and then 3-5µl were loaded on 5% (v/v) native 0.5X TBE PAGE gel with 5 mM MgCl_2,_ which was prerun at 8W for ∼1hr in 4°C. Gel was running in 0.5X TBE buffer with 5 mM MgCl_2_ for ∼2.5hr at 8W in 4°C. Gel was dried and phosphorous screen was exposed with the gel overnight. Gel was imaged on Typhoon and quantified using ImageQuant software. Fraction of promoter sequence bound was calculated by dividing the signal of bound DNA fraction by the sum of bound and unbound signals in each lane. Signal was considered to come from the bound state if it was above the level of unbound state determined by the 0nM GABP point. Signal was considered fully bound for the highest shifted bands and the signal between unbound and fully bound was considered intermediate bound state. Replicates were obtained by preparing one 2 µM heterodimer stocks which was further serially diluted to create 15 new stocks and equal volume of protein from each stock was added to three separate sets of tubes for each DNA probe tested.

### Mammalian cell culture

All mammalian cell cultures were maintained in a 37°C incubator at 5% CO_2_. HEK293T human kidney cells (293FT; Thermo Fisher Scientific, #R70007; RRID:CVCL_6911), NHA-PC5 normal human astrocyte cells (a kind gift from Russell Pieper, UCSF), and derivatives thereof were grown in Dulbecco’s Modified Eagle Medium (DMEM; Corning Cellgro, #10-013-CV) supplemented with 10% fetal bovine serum (FBS; Seradigm #1500-500), and 100 Units/ml penicillin and 100 µg/ml streptomycin (100-Pen-Strep; Gibco, #15140-122). U-251 human glioblastoma cells (Sigma-Aldrich, #09063001; RRID:CVCL_0021), LN-229 human glioblastoma cells (American Type Culture Collection (ATCC), #CRL-2611; RRID:CVCL_0393), T98G human glioblastoma cells (ATCC, #CRL-1690; RRID:CVCL_0556), LN-18 human glioblastoma cells (ATCC, #CRL-2610; RRID:CVCL_0392), and derivatives thereof were cultured in Dulbecco’s Modified Eagle Medium/Nutrient Mixture F-12 (DMEM/F-12; Gibco, #11320-033 or Corning Cellgro, #10-090-CV) supplemented with 10% FBS and 100-Pen-Strep. HAP1 cells (a kind gift from Jan Carette, Stanford University) and derivatives thereof were grown in Iscove’s Modified Dulbecco’s Medium (IMDM; Gibco, #12440-053 or HyClone, #SH30228.01) supplemented with 10% FBS and 100-Pen-Strep. HAP1 cells had been derived from the near-haploid chronic myeloid leukemia cell line KBM7^64^.

U-251, LN-229, T98G, LN-18, HEK293T, and HAP1 cells were authenticated using short tandem repeat DNA profiling (STR profiling; UC Berkeley Cell Culture/DNA Sequencing facility). STR profiling was carried out by PCR amplification of nine STR loci plus amelogenin (GenePrint 10 System; Promega, #B9510), fragment analysis (3730XL DNA Analyzer; Applied Biosystems), comprehensive data analysis (GeneMapper software; Applied Biosystems), and final verification using supplier databases including ATCC and Deutsche Sammlung von Mikroorganismen und Zellkulturen (DSMZ).

U-251, LN-229, T98G, LN-18, NHA-PC5, HEK293T, and HAP1 cells were initially tested for absence of mycoplasma contamination (UC Berkeley Cell Culture facility) by fluorescence microscopy of methanol fixed and Hoechst 33258 (Polysciences, #09460) stained samples. Lack of mycoplasma contamination was confirmed every 6 months using MycoAlert mycoplasma detection kit (Lonza, #LT07-418).

### Establishment of a puromycin-sensitive normal human astrocyte cell line

The normal human astrocyte cell line NHA-PC5 was previously described^45^, and is puromycin resistant. For our experiments, we established a puromycin-sensitive monoclonal derivative, termed NHA-S2, through CRISPR editing of the previously inserted puromycin resistance cassette. In brief, NHA-PC5 cells were transiently transfected in 6-well plates with a pair of plasmid vectors (pCF120), each encoding Cas9 and a guide RNA targeting the puromycin resistance marker (sgPuro5 and sgPuro6, Table S1), or a negative control. Transfections were carried out using Lipofectamine 3000 (Thermo Fisher Scientific) according to the manufacturer’s procedures (500 ng vector 1, 500 ng vector 2). For construction of pCF120 see corresponding section (Plasmid and lentiviral vectors). Starting at 24 h post-transfection, NHA-PC5 cells were selected on hygromycin B (500 µg/ml; Thermo Fisher Scientific, #10687010) for four days.

Subsequently, the selected cells were seeded using limiting dilution, and 16 clones picked 17 days later. Clones were further expanded and tested for puromycin (1.0 µg/ml; InvivoGen, #ant-pr-1) sensitivity. All 11 out of 11 edited NHA-PC5 clones that grew well were puromycin sensitive. Five of these clones, NHA-S1/S2/S3/S7/S9 (S for puromycin sensitive), were further expanded and tested for hygromycin B (500 µg/ml) sensitivity to ensure that the plasmid vectors (pCF120) used for editing did not integrate into the genome. NHA-S3 were hygromycin B resistant, NHA-S1 were partially resistant, and NHA-S2/S7/S9 showed sensitivity. From the latter three, NHA-S2 were chosen as the cell line for further experiments based on morphology and cell growth. NHA-S2 were authenticated using short tandem repeat DNA profiling (STR profiling; UC Berkeley Cell Culture/DNA Sequencing facility) to be derived from NHA-PC5. NHA-PC5 and NHA-S2 were stored in freezing medium consisting of 5% DMSO, 40% FBS, and 55% growth medium.

### Sequencing of *TERT* promoter mutation status

To sequence *TERT* promoter mutation status, genomic DNA was extracted from cell lines using QuickExtract DNA Extraction Solution (Epicentre, #QE09050). Specifically, samples were incubated at 65°C for 20 minutes followed by 20 minutes at 98°C. The *TERT* promoter locus was amplified with the primers oCF1039_TERTp-fw2 (CAGGGCCTCCACATCATGG) and oCF1038_TERTp-rev1 (CCTCGCGGTAGTGGCTG), yielding a 532 bp product. PCRs were carried out using Q5 High-Fidelity DNA polymerase (New England Biolabs, #M0491) with an annealing temperature of 70°C, 20 seconds elongation time, 32 PCR cycles, and standard Q5 buffer with 10% High-GC Enhancer due to elevated G/C content of this locus. Sanger sequencing of the *TERT* promoter locus was carried out, after gel purification of the PCR bands, using the primers oCF1039_TERTp-fw2 and oCF1038_TERTp-rev1. Sequences were analyzed using SnapGene software (GSL Biotech).

### Plasmid and lentiviral vectors

To generate monoclonal knockout cell lines, sgRNAs and Cas9 were expressed from either the pCF120 or pCF123 vectors. In brief, the plasmid vector pCF120, expressing a U6 promoter driven sgRNA and an EF1-alpha short (EFS) promoter driven Cas9-P2A-Hygro cassette, was derived from pX459^65^ by replacing the chicken beta-actin promoter with an EFS promoter from pCF204^66^, swapping the T2A-Puro cassette with a P2A-Hygro cassette, and exchanging the guide RNA scaffold for a more efficient variant^67^. The plasmid vector pCF123, expressing a U6 driven sgRNA and an EFS driven Cas9-T2A-Puro cassette, was generated based on pX459 by replacing the chicken beta-actin promoter with an EFS promoter and exchanging the guide RNA scaffold for a more efficient variant^67^.

For CRISPR-Cas9-mediated competitive proliferation assays, single-guide RNAs (sgRNAs) were expressed from the previously established pCF221 lentiviral vector and Cas9 from the pCF226 lentiviral vector^66^. For low-level stable overexpression of *GABPB2* (NCBI gene ID: 126626) and mTagBFP2 (negative control), we first modified the pCF525-mTagBFP2 lentiviral vector^68^, encoding an EF1a-Hygro-P2A-mTagBFP2 cassette, by replacing the strong EF1-alpha promoter with a weak EF1-alpha short (EFS) promoter. This new vector, EFS-Hygro-P2A-mTagBFP2, was named pCF554. Next, we cloned the coding sequence (CDS) of homo sapiens GA binding protein transcription factor subunit beta 2 (GABPB2), transcript variant 1, mRNA (NCBI reference sequence: NM_144618.2) into pCF554 by replacing mTagBFP2, yielding lentiviral vector pCF553. The *GABPB2* CDS was synthesized as gBlocks (Integrated DNA Technologies).

For TERT rescue experiments, we stably overexpressed BFP (control) or TERT (rescue) using commercially available lentiviral constructs. Specifically, 10-30% confluent U-251 cells were transduced at low MOI in 12-well plates with the lentivirus EF1a-BFP_Rsv-Bsd (5 µl; GenTarget, #LVP365) or EF1a-hTERT_Rsv-Bsd (50 µl; GenTarget, #LVP1131-Bsd). The EF1a-hTERT_Rsv-Bsd (GenTarget, #LVP1131-Bsd) vector expresses human TERT based on NM_198253.2, but does not include any 3’ UTR sequence of the natural mRNA and the stop codon was switched from the native TGA to TAG. At 24 h post-transduction, transduced cells were selected on Blasticidin S (10 µg/ml; Gibco, #A1113903) for 7 days. For doxycycline-inducible miR-E shRNA expression, both in cell culture and *in vivo*, we used the lentiviral vector LT3GEPIR^36^. LT3GEPIR provides a GFP marker reporting shRNA expression. Full sequences of all vectors are provided (Table S1). All vectors were cloned and/or sequence verified using standard molecular biology techniques, enzymes from New England Biolabs (NEB), and oligonucleotides from Integrated DNA Technologies (IDT).

### Lentiviral transduction

Lentiviral particles were produced in HEK293T cells using polyethylenimine (PEI; Polysciences #23966) based transfection of plasmids, as previously described^66^. In brief, lentiviral vectors were co-transfected with the lentiviral packaging plasmid psPAX2 (Addgene #12260) and the VSV-G envelope plasmid pMD2.G (Addgene, #12259). Transfection reactions were assembled in reduced serum media (Opti-MEM; Gibco, #31985-070). For lentiviral particle production on 6-well plates, 1 µg lentiviral vector, 0.5 µg psPAX2 and 0.25 µg pMD2.G were mixed in 0.4 ml Opti-MEM, followed by addition of 5.25 µg PEI. After 20-30 min incubation at room temperature, the transfection reactions were dispersed over the HEK293T cells. Media was changed 12-14 h post-transfection, and virus harvested at 42-48 h post-transfection. Viral supernatants were filtered using 0.45 µm polyethersulfone (PES) membrane filters, diluted in cell culture media as appropriate in order to obtain similar levels of viral expression between cell lines and conditions, and added to target cells. Polybrene (5 µg/ml; Sigma-Aldrich) was supplemented to enhance transduction efficiency, if necessary.

### Design of sgRNAs for CRISPR-Cas9 genome editing

Standard sgRNA sequences were either designed manually or using GuideScan^69^. To generate specific genomic excisions, rather than indels, pairs of sgRNAs were designed and used in tandem. This strategy was applied to remove sequences specific to the long isoform of *GABPB1L*, while not affecting the short isoform. All sgRNAs were designed with a G preceding the 20 nucleotide spacer for better expression from U6 promoters. All sgRNA sequences are shown (Table S1), and were cloned into the pCF120 and pCF123 vector using BbsI restriction sites and enzymes (New England Biolabs) or into the pCF221 vector using Esp3I restriction sites and enzymes (New England Biolabs). Control sequencing of sgRNAs was carried out with the primer oCF112_U6seq2-fw (ACGATACAAGGCTGTTAGAGAG). Oligonucleotides for sgRNA cloning were purchased from IDT.

### CRISPR-Cas9 competitive proliferation assay

CRISPR-Cas9 competitive proliferation assays were used to assess whether Cas9-mediated editing of genes of interest leads to a change in proliferation speed compared to unedited cells. First, cell lines (U-251) were stably transduced with a lentiviral vector expressing Cas9 (pCF226) and selected on puromycin (1.0-2.0 µg/ml), as previously described^66^. Subsequently, Cas9 expressing cell lines were further stably transduced with pairs of lentiviral vectors (pCF221) expressing various mCherry-tagged sgRNAs. At day two post-transduction, sgRNA expressing populations were mixed approximately 80:20 with parental cells and the fraction of mCherry-positive cells was quantified over time by flow cytometry (Attune NxT flow cytometer, Thermo Fisher Scientific).

### Design of shRNAs for reversible target inhibition

MicroRNA-embedded miR-E shRNA sequences were predicted using SplashRNA^38^ and cloned into LT3GEPIR^36^, an all-in-one doxycycline-inducible miR-E shRNA expression vector. The shRNA numbers refer to the position of the shRNA, specifically the 3’ nucleotide of the guide strand, on the target transcript^38, 70^. All shRNA sequences are shown (Table S1), and were cloned as previously described^70^. Oligonucleotides for shRNA cloning were purchased from IDT.

### Generation of isogenic GABPB1L knockout cell lines

To generate isogenic monoclonal knockout cell lines of the long isoform of GABPB1 (NCBI gene ID: 2553), pairs of sgRNAs flanking the long isoform-specific exon 9 of *GABPB1* were used. In brief, the sgRNAs sgGB.2, sgGB.3, sgGB.4, sgGB.12, sgGB.13, and sgGB.14 as well as sgPTEN.3 and sgPTEN.12 (negative controls) were cloned into the pCF123 plasmid vector (Table S1). To test genome editing and locus excision efficiency, every possible combination of a 5’ flanking (intron) and a 3’ flanking (3’UTR) sgRNA were tested alonside controls by transient transfection into both HEK293T and U-251 cells. Transfections were carried out in 12-well plates using Lipofectamine 3000 (Thermo Fisher Scientific) according to the manufacturer’s procedures (250 ng vector 1, 250 ng vector 2). Starting at 24 h post-transfection, HEK293T and U-251 cells were selected on puromycin (HEK293T: 1.0 µg/ml, U-251: 0.5 µg/ml) for two days. Subsequently, cells were expanded and genomic DNA extracted (QuickExtract DNA Extraction Solution; Epicentre, #QE09050), as described above, for genotyping analysis. Genotyping, as described below, determined sgGB.2-12 and sgGB.3-14 to be the two most efficient guide pairs (Fig. S2E).

GABPB1L locus editing was assessed by genotyping with the primers oCF862_GAL-fw1 (AAAAGTACAGGTGCCCAGTTTG) and oCF863_GAL-rev2 (GCCTAACCAACAACGATCAC), yielding an 808 bp band in wild-type cells. Genotyping PCRs were carried out using Q5 High-Fidelity DNA polymerase (New England Biolabs, #M0491) with an annealing temperature of 65°C, 20 seconds elongation time, and 32 PCR cycles. Where mentioned, Sanger sequencing of the GABPB1L locus and TIDE analysis^71^ were also carried out using the primers oCF862_GAL-fw1 and oCF863_GAL-rev2. Perfect GABPB1L editing with the sgRNA pair sgGB.2-12 yields a 472 bp genotyping band, and with the sgGB.3-14 pair a 387 bp band. To establish isogenic monoclonal GABPB1L knockout cell lines, TERT promoter wild-type (HEK293T, HAP1, NHA-S2, LN-18) and TERT promoter mutant (U-251, LN-229, T98G) cell lines were edited with pCF123-sgGB.2-12 and pCF123-sgGB.3-14 pairs in 12-well (250 ng vector 1, 250 ng vector 2) or 6-well (500 ng vector 1, 500 ng vector 2) plates, as described above. After selection on puromycin (0.5-3.0 µg/ml), the selected cells were seeded using limiting dilution, and clones picked approximately 10-15 days later. Clones were then expanded, genotyped (Fig. S3), and further cultured for secondary analysis and storage. A set of clones from each cell line were selected for further analysis based on clean sequencing of the edited locus, and normal morphology and growth. A mix of clones generated using each of the 2 different pairs of sgRNAs were selected for each cell line. Selected clones were genotyped a second time (Fig. 2C,D and Fig. S4) and authenticated using short tandem repeat DNA profiling (STR profiling; UC Berkeley Cell Culture/DNA Sequencing facility) to ensure that monoclonal lines are derived from the intended parental cell line. Specifically, the following clones were authenticated by STR profiling: LN-229 C6, C10, C11, C19, C24, C26, C28, C30, LN-18 C4, C8, C11, C14, C21, C23, C24, C32, T98G C3, C4, C5, C9, C13, C20, and NHA-S2 C1, C4, C14, C23, C24, C29, C32. All secondary analyses performed on isogenic clones were carried out using aliquots of cells frozen 25-35 days post-editing, in order to minimize possible long-term effects on cell state, telomere attrition, or viability resulting from GABPB1L editing.

### Flow cytometry and fluorescence microscopy

Percentages of fluorophore-positive cells were quantified by flow cytometry (Attune NxT flow cytometer, Thermo Fisher Scientific), regularly acquiring 10,000-30,000 events per sample. Phase contrast imaging and fluorescence microscopy was carried out following standard procedures (EVOS FL Cell Imaging System, Thermo Fisher Scientific), routinely at least 48 h post-transfection or post-transduction of target cells with fluorophore expressing constructs.

### RT-qPCR

Cells were seeded in 96 well plates in a volume of 100ul, at a concentration of 50 cells/ul. When cells reached ∼80% confluency (3-4 days post-seeding), RNA isolation and cDNA generation was performed using the Cells-to-CT kit (Invitrogen, #4402954), per the manufacturer’s instructions. Real-time quantitative PCR was performed using the POWER SYBR Green PCR Master Mix (Applied Biosystems, #4367659) to measure expression levels of *GUSB, TERT*, *GABPB1L*, total *GABPB1*, *GABPB1S*, *GABPB2*, *RPA1,* and *MGMT*. Each sample was measured in technical and biological triplicate on a QuantStudio 5 Real-Time PCR machine (Thermo Fisher Scientific). Manual inspection of melting curves was performed to confirm PCR specificity. Relative expression levels of biological triplicates were quantified using the deltaCT method against *GUSB*. Experiments were performed three times with separate RNA isolations for final quantification. Primer sequences are available in Table S1.

### Immunoblotting

Cells were washed two times in cold PBS and harvested on ice with M-PER Mammalian Protein Extraction Reagent (Thermo Scientific, #78501), plus freshly added Turbonuclease (Sigma Aldrich, #T4330), and Halt Protease and Phosphatase Inhibitor Cocktail (Thermo Scientific, #1861282). Protein concentration was determined by Bicinchoninic acid (BCA) assay. Equal protein amounts were resolved on SDS-PAGE gels and electrotransferred to a PVDF membrane. Membranes were blocked with 5% BSA in Tris-buffered saline tween 20 (TBST 0.1%) or 5% milk in Tris-buffered saline tween 20 (TBST 0.1%) for 1 hour, and then probed with primary antibodies towards GABPB1 (Proteintech, #12597-1-AP), yH2AX (Cell Signaling, #80312S), or GAPDH (Millipore, #CB1001). Each immunoblot was performed a minimum of three times with separate protein samples.

### Telomerase Repeated Amplification Protocol (TRAP)

Cy5-based TRAP assays were performed as previously described^72^. Briefly, 150,000 cells per condition were collected via trypsinization and resuspended in NP-40 lysis buffer (10mM Tris-HCl pH 8.0, 1mM MgCl_2_, 1mM EDTA, 1% NP-40, 0.25mM sodium deoxycholate, 10% glycerol, 150mM NaCl, 5mM 2-mercaptoethanol, 0.1mM AEBSF) at a concentration of 2,500 cells/ul to maintain telomerase activity. 1 ul of lysate (or 1 ul of NP-40 lysis buffer for the negative control) was added to a PCR master mix containing a Cy5-labeled telomeric substrate primer, an internal standard control, and an extension primer mix, at the described ratios^72^ The following PCR protocol was used to amplify Cy5 telomeric primer extension products: 1) 25°C for 40 min; 2) 95°C for 5 min; 3) 27 cycles of 95°C for 30 sec, 52°C for 30 sec, 72°C for 45 sec; 4) 72°C for 10 min. Post-PCR samples were resolved on a polyacrylamide gel, and visualized using a BioRad Chemidoc Imager on the Cy5 setting. Data was visualized and analyzed using BioRad Image Lab software 6.0 and ImageJ with n=3-4 biological replicates.

### Orthotopic xenografts and bioluminescence imaging

Animal procedures conformed to care guidelines approved by the UCSF Institutional Animal Care and Use Committee (IACUC protocol numbers: AN179550, AN17599, AN179565). U-251 and LN-229 cell lines were stably transduced with Firefly Luciferase Lentifect Purified Lentiviral Particles (Genecopoiea, #LPP-FLUC-Lv105) at an MOI of 3. All cell lines were verified to be stably expressing luciferase at similar levels *in vitro* 48 hours prior to xenograft. Either 4×10^5^ U-251 cells or 1×10^5^ LN-229 cells were injected in a 4ul total volume into the right frontal cortex (ML:1.5mm, AP:1mm, DV:3.5mm) of 6-7 week old female athymic *nu/nu* mice (inducible shRNA TMZ combination experiment: The Jackson Laboratory, RRID:IMSR_JAX:002019; all other experiments: Envigo, Hsd:Athymic Nude-Foxn1^nu^, #069). Sample sizes were estimated based on both the standard in the field, as well as power calculations using previously observed effect sizes published in experiments using similar techniques. Later experiments were re-estimated based on observed effect sizes in our own work. Animals were excluded from analysis if surgical complications arose immediately (within 48 hours) post-surgery. Approximate tumor size was monitored via bioluminescence imaging (Xenogen IVIS Spectrum Imaging System) 1-2x per week following intraperitoneal injection of luciferin reagent (GOLDBIO, #LUCK-1G) at 150mg/kg. Animals were weighed 3x per week, and once daily after initial weight loss was observed. General behavior and symptomology for all mice were recorded daily. Humane endpoints for sacrifice were hunched posture, >15% weight loss from maximum recorded weight, or neurologic symptoms, and were verified by approved UCSF veterinary technicians.

### *In vivo* doxycycline administration and analysis

Mice bearing xenografts of U-251 cells expressing doxycycline-inducible shRNAs were placed on either control or doxycycline chow at a dosage of 625 mg/kg (Dox Diet; Bio-Serv, #F5829) seven days post xenograft, when bioluminescence readings were in the 1×10^6^ (radiance) range. Mice were placed into an experimental condition post-xenograft (control versus doxycycline chow) via randomization based on tumor volume, as determined by bioluminescence imaging. Mice were maintained on appropriate chow until humane endpoint. To analyze protein levels or induced GFP expression in U-251 orthotopic xenografts, tumors were dissected from the brain *post mortem*, and dissociated into a single cell suspension through manual trituration followed by passage through a 100um cell strainer. A portion of the cell suspension was washed in cold PBS and subjected to lysis for immunoblot analysis, as previously described. For analysis of GFP expression, cells were washed 2x in FACS buffer (PBS + 2% fetal bovine serum + 0.05% Sodium Azide), then resuspended in FACS buffer containing 2 ug/ml APC/Cy7 anti-human HLA-A,B,C (BioLegend, #311425). Cells were incubated with the antibody on ice for 20 minutes in the dark, then washed 2x with FACS buffer, and resuspended in FACS buffer for flow cytometry analysis. GFP and Far Red (APC-Cy7) fluorescence were measured by flow cytometry (Attune NxT, Thermo Fisher Scientific), routinely acquiring 10,000 events per sample. Percent of GFP positive tumor cells was calculated as the percentage of Far Red positive cells (human cells) that were also GFP positive.

### Temozolomide administration *in vitro*

U-251 and LN-229 cells were seeded in 24 (600ul/well) well plates at a density of 500 cells/ul. For cells transduced with doxycycline-inducible shRNA constructs, cells were maintained on doxycycline at 1 ug/ml for 4 days prior to plating and through TMZ treatment paradigm. 24 hours post-seeding, cells were treated with the described concentrations of TMZ (Tocris, #85622-93-1), or a DMSO control, in a total volume of 2ul/ml, for 3 hours. TMZ drug was then removed. For yH2AX immunoblots, cells were harvested 20 hours post TMZ treatment. At the end of the 20 hour incubation, cells were harvested for analysis by western blot as described above and analyzed by quantification using Image J. For cell cycle analysis, cells in 24 well plates were harvested 72 hours post-TMZ treatment and fixed in 70% ethanol at 4°C overnight. Fixed cell pellets were washed 2x with cold PBS, resuspended in 300 ul FxCycle PI/RNAse solution (ThermoFisher Scientific, #F10797), and incubated at room temperature in the dark for 15 minutes. Samples were then analyzed by flow cytometry, using the Attune NxT (Thermo Fisher Scientific) flow cytometer, routinely acquiring 10,000 events per sample. Experiments were done in biological triplicate.

### Temozolomide administration *in vivo*

Mice were placed into an experimental condition post-xenograft (temozolomide versus vehicle) via randomization based on tumor volume, as determined by bioluminescence imaging. Mice assigned to the temozolomide condition were given *per os* (p.o.) doses of Temodar temozolomide capsules (NDC, #43975-257-05) at either 50mg/kg for 10 days total (5 days on, 2 days off, for a period of 12 days; unaltered or GABPB1L-edited U-251 xenografts) or 10mg/kg for 5 days (5 consecutive days; LN-229 xenografts and shRNA-expressing U-251 xenografts). In the case of Temodar administration to mice bearing doxycycline-inducible shRNA transformed U-251 xenografts, mice were placed on doxycycline chow for 5 days prior to Temodar treatment. Temodar was resuspended in Ora-Plus oral suspension vehicle at either 5mg/ml (for 50mg/kg) or 1mg/ml (for 10mg/kg), to be administered at 10ml/kg (0.2ml per 20g mouse). Mice assigned to the vehicle condition were given an equivalent volume of Ora-Plus without drug. Both Temodar and vehicle solutions were administered to mice by an experimenter blind to the xenograft condition. Simple additive effects of TMZ combined with GABPB1L knockout were calculated as 1+((M_y_/M_x_-1)+(M_z_/M_x_-1)), where M_x_=median survival of control vehicle condition, M_y_=median survival of control TMZ condition, and M_z_=median survival of GABPB1L or TERT knockout vehicle condition.

### Imaging and Analysis

For immunofluorescence-based analysis of GFP, Ki-67, and GFAP expression in inducible shRNA-expressing U-251 tumors treated with vehicle or TMZ, animals were euthanized 30 days post-xenograft to compare across conditions at identical timepoints. Animals were perfused, and brains fixed in 4% PFA for 24 hours. Brains were then washed in PBS, and soaked in 30% sucrose prior to freezing in OCT blocks for cryosectioning. Slides were washed in PBS to remove OCT, blocked in 5% BSA + 0.3% Triton-X in PBS for 1 hour, then incubated with primary antibodies towards Ki-67 (Thermo Fisher, #PA5-19462) or GFAP (Abcam, #ab4674-50) overnight, followed by secondary antibodies for 1 hour (Alexa Fluor 647, Abcam #ab150171 or Alexa Fluro 555, Thermo Fisher #A27039). Slides were incubated with Hoechst stain for 10 minutes prior to mounting. GFP immunofluorescence was detectable without the need for antibody staining. Imaging was performed using a Keyence digital microscope, by a researcher blinded to experimental condition. 2X representative images were taken of the entire tumor field for each of 3 mice per condition. 40X images near the tumor edges were obtained for analysis of Ki-67 expression, since Ki-67 expression is highest at the tumor edge, and is used as a pathology-based marker for tumor grade. Ki67 image quantification was carried out using similar methods to previously published work^73–75^. Briefly, DAPI, Ki67 and GFP channels of each image were imported into MATLAB 2020a. The Ki67 channel was thresholded at a defined value of 40. This value was determined by manually thresholding several raw images to ensure that Ki67 cells were accurately classified. DAPI and GFP channels were subjected to local contrast normalization by dividing each image by the local intensity of its neighboring pixels (see code for details). The contrast normalized DAPI images were then thresholded and segmented using a seeded watershed algorithm implemented in the DIPimage toolbox 2.9. The overlap between segmented DAPI, GFP and Ki67 was used to categorize each nuclei as positive for each channel. Full MATLAB script available at: https://github.com/mabdullahsyed/amen-2020. (Note: The DIPimage toolbox is required to run this script). Quantification was performed using 5 fields per tumor and 3 tumors per condition.

### siRNA knockdown

Non-targeting (#D-001206-13-20), GABPA (#M-011662-01), and GABPB2 (#M-016074-00-0005) directed siGENOME SMARTsiRNA pools were purchased from Dharmacon. LN-229 cells were seeded in a 100ul volume in 96 well plates at a concentration of 500 cells/ul. 24 hours post-seeding, cells were transfected using 50 nM siRNA and 0.1 ul Dharmafect I reagent (Dharmacon, #T-2001-01). At 72 hours post-transfection, cells were lysed and harvested for qPCR analysis, as described above.

### Statistical analysis

Statistical analyses were performed using GraphPad Prism 9 software. Two-tailed, unpaired student’s *t* tests were used with alpha = 0.01 or alpha = 0.05, as specified. Statistical significance between Kaplan Meier survival curves was performed using a Kaplan Meier log-rank test with alpha = 0.01 or alpha = 0.05, as specified. The Wilcox signed-rank test was used for comparison of GABPB1 and GABPB2 expression levels from TCGA RNAseq datasets. Statistical analysis for qPCR data was performed among specific samples, rather than relative to wild-type control, due to normalization on wild-type control using the deltaCT method. Quantified data represent mean +/- standard deviation or mean +/- SEM, as referenced in figure legends. Values of ‘n’ are listed in relevant methods and/or figure legends.

## Acknowledgements

We thank R. Pieper (UCSF) for generously sharing the immortalized astrocyte line, NHA-PC5, and for consultation on experiments involving temozolomide. We thank the UCSF Preclinical Therapeutics Core and the UCSF Preclinical Animal Core for their assistance with animal experiments and maintenance of imaging equipment. We thank Mary West and the University of California, Berkeley CIRM/QB3 Shared Stem Cell Facility/High-Throughput Screening Facility for support. We thank the Gladstone Institute Histology and Microscopy core for assistance with cryosectioning and support. We thank Abdullah Syed for assistance with image analysis. This work was supported by NIH grant NCI F32CA228365 (A.M. Amen), NIH K99/R00 Pathway to Independence Award NIGMS K99GM118909 and R00GM118909 (C. Fellmann), NCI fellowship F99 CA222987 (A. Mancini), NCI fellowship F31 CA243187 (A.M. McKinney), a generous gift from Mitchel Berger and the Fishgold Hurwitt Brain Tumor Research Fund (C. Fellmann and J.A. Doudna), NIH grant NCI P50CA097257 (J.F. Costello and J.A. Doudna), funding from the GBM Precision Medicine Project (J.F. Costello and J.A. Doudna), a generous gift from the Dabbiere family (J.F. Costello), and a generous gift from the Hana Jabsheh Research Initiative (J.F. Costello). J.A. Doudna is an investigator of the Howard Hughes Medical Institute. We also thank all Doudna and Costello lab members for insightful discussions.

## Author Contributions

A.M.A., C.F., J.A.D, and J.F.C. conceived the project. A.M.A. and C.F. designed, planned, and carried out experiments, and analyzed and interpreted data. K.M.S, G.J.K, and J.E.P. conceived, designed, and carried out protein purification and DNA binding assays. S.M.R. and R.J.L. helped plan and carry out experiments. A.M.M. and A.M. helped plan experiments and interpret data. J.A.D. and J.F.C supervised all experiments. A.M.A., C.F., J.A.D, and J.F.C. wrote the manuscript with input from all authors.

## Supplementary Figure Legends

**Supplementary Fig. 1.**
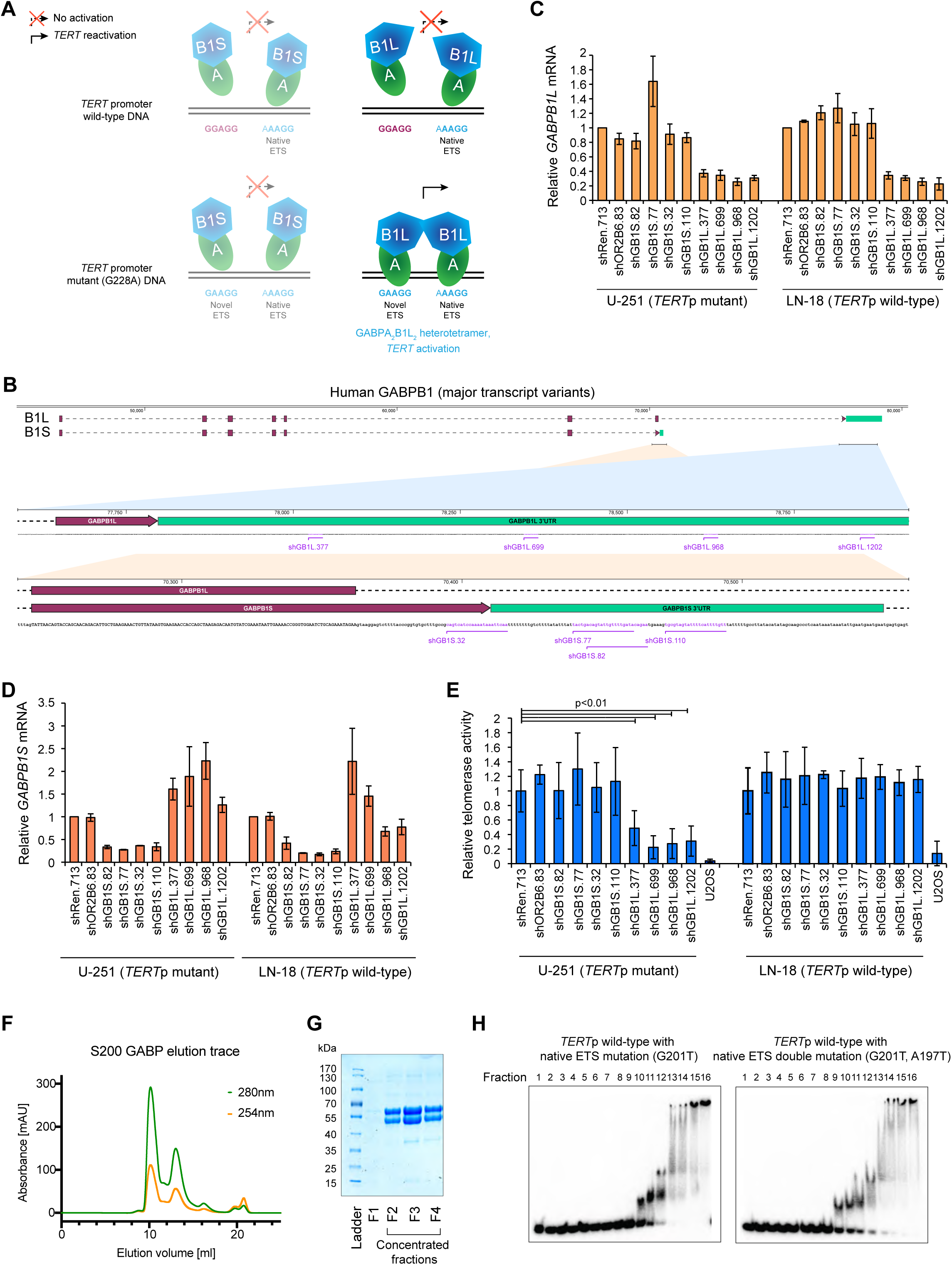
Regulation of TERT by GABPB1 isoforms. **A**, Schematic of GABP complex binding to the wild-type versus mutant *TERT* promoter. The mutant promoter contains two overlapping native ETS family transcription factor binding sites that are hypothesized to recruit a GABPB1L-containing heterotetramer complex, GABPBA_2_B1L_2_. **B**, Schematic of GABPB1 short and long isoform-specific shRNAs. Shown are the major human *GABPB1* transcript variants, encoding either a short (*GABPB1S*) or long (*GABPB1L*) isoform. Red rectangles represent the coding sequence (CDS), green rectangles the 3’ untranslated region (3’UTR). For clarity, the first exon containing the 5’ untranslated region (5’UTR) is not shown. GABPB1S and GABPB1L targeting shRNAs (shGB1S, shGB1L) are highlighted in purple. **C**,**D**, GABPB1L (**C**) and GABPB1S (**D**) mRNA expression levels measured via qRT-PCR in U-251 and LN-18 cells expressing doxycycline-induced shRNAs targeting GABPB1S (shGB1S.82, shGB1S.77, shGB1S.32, shGB1S.110) and GABPB1L (shGB1L.377, shGB1L.699, shGB1L.968, shGB1L.1202) compared to negative control (olfactory receptor OR2B6, shOR2B6.83) and non-targeting (renilla luciferase, shRen.713) shRNAs. Cells were incubated with doxycycline for 6 days prior to harvest. **E,** Quantification of telomerase activity, measured via TRAP assay, in U-251 and LN-18 cell lines from (**C**,**D**), and a control cell line (U2OS) lacking *TERT* expression. Telomerase banding intensity was normalized to internal control band and plotted relative to shRen.713. p: p-value (unpaired, two-tailed student’s t-test). Data represent mean +/- standard deviation. **F**,**G**, Representative superdex column trace (**F**) and acrylamide gel (**G**) for purification of GABPA-B1L heterodimers. Observed molecular weights of GABPA ≈ 55 kDa, GABPB1L ≈ 50 kDa. **H**, Representative gels of EMSAs assessing binding affinity of GABPA-B1L heterodimers to wild-type *TERT* promoter sequences lacking native ETS binding sites (G201T: native ETS single mutant; G201T, A197T: native ETS double mutant).

**Supplementary Fig. 2.**
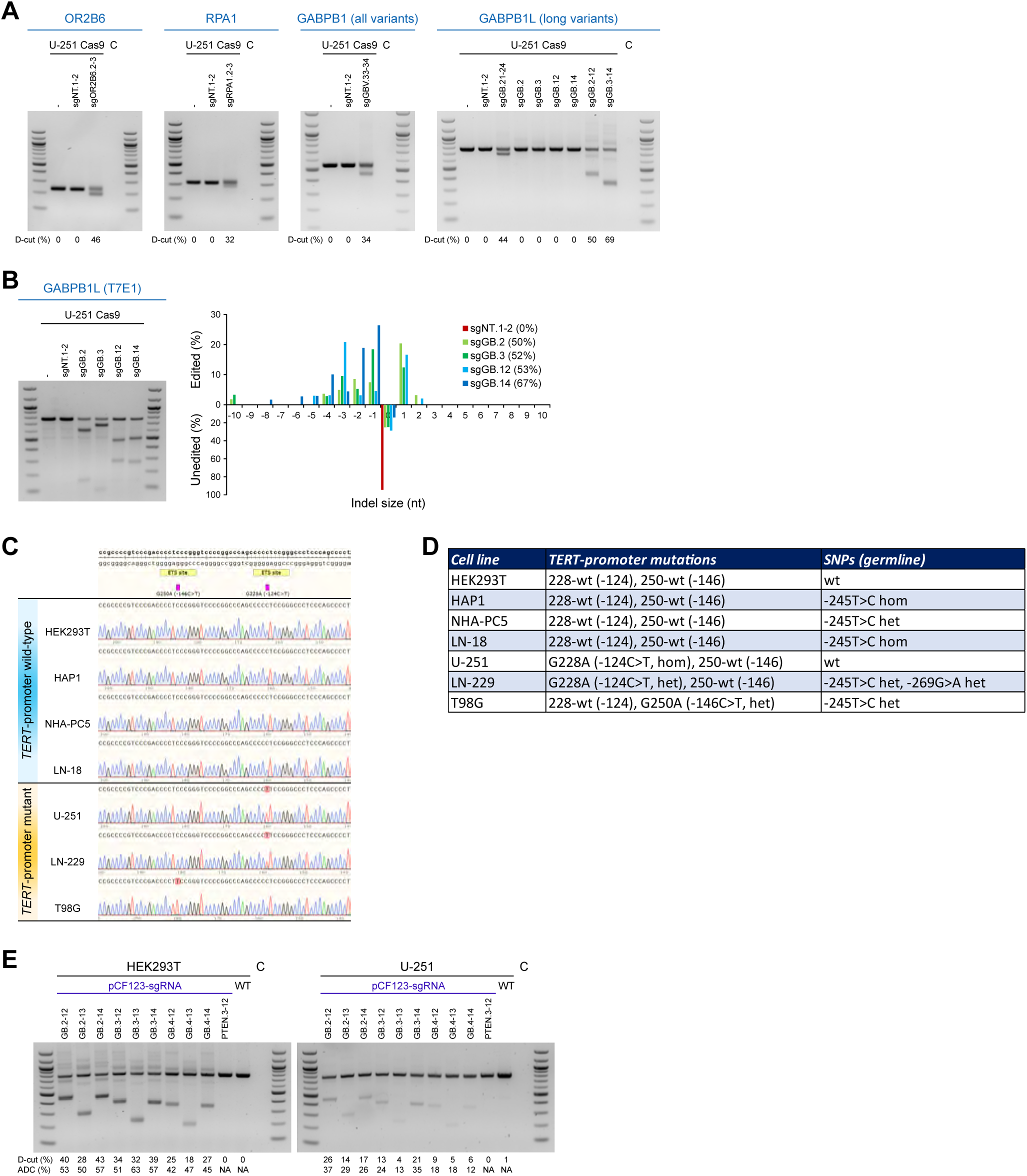
Editing of *GABPB1L* in *TERT* promoter wild-type and mutant cell lines. **A**, Genotyping gels showing editing efficiency of sgRNA editing. U-251 cells stably expressing Cas9 from the lentiviral vector pCF226 were transduced with pairs of lentiviral vectors (pCF221) expressing the indicated sgRNAs. Genomic DNA was harvested at day 4 post-transduction. Genotyping PCRs were run at the indicated loci (Table S1) and locus excision efficiency quantified (ImageJ). D-cut: double-cut. C: PCR no template control. **B**, Quantification of sgRNA editing efficiency in U-251 GBM cells. Indel generation efficiency of guide RNAs was assessed by T7 endonuclease I (T7E1) assay and quantified by deconvolution of sequencing reads (TIDE). **C**, Sequencing of *TERT* promoter mutation status. Representative sequencing chromatograms of a select region of the *TERT* promoter for the indicated cell lines are shown. **D**, Summary of *TERT* promoter mutation status and nearby single nucleotide polymorphisms (SNPs) in the indicated cell lines. wt: wild-type. **E**, Analysis of different sgRNA pairs targeting *GABPB1L* intron 8 and 3’ UTR to excise exon 9 of *GABPB1L*. PCR-based genotyping analysis of pooled editing of a *TERT* promoter wild-type control cell line (HEK293T) and a *TERT* promoter mutant GBM cell line (U-251) through transient transfection of vectors (pCF123) expressing Cas9 and the indicated pairs of sgRNAs, followed by puromycin selection. C: PCR water control. wt: parental cell line wild-type control. D-cut: relative double-cut band intensity compared to full-length, quantified by densitometry (ImageJ). ADC: D-cut normalized to PCR fragment length.

**Supplementary Fig. 3.**
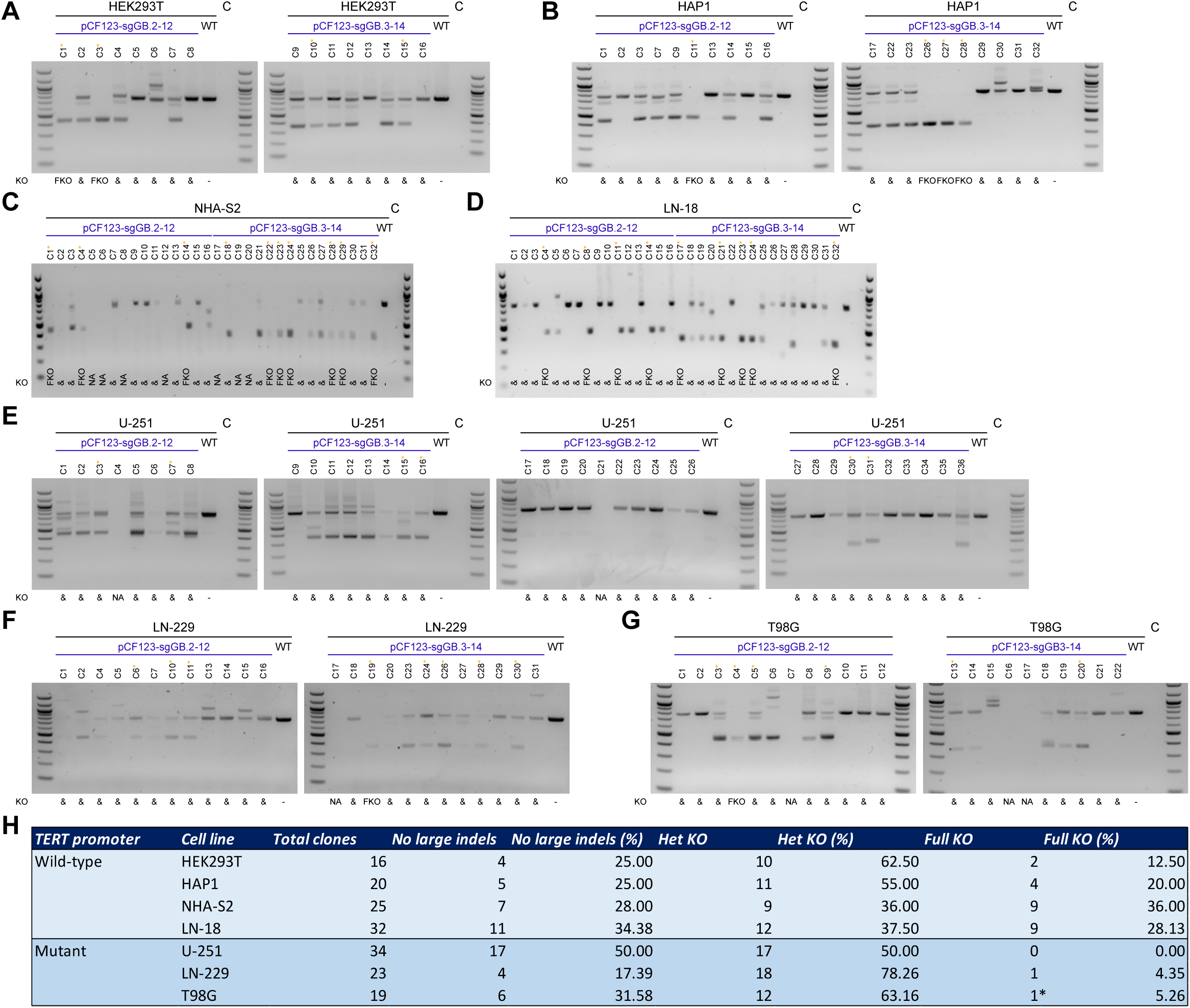
Generation of monoclonal GABPB1L full and heterozygous knockouts in *TERT* promoter wild-type and mutant cell lines. **A-G**, Establishment of *GABPB1L* edited isogenic monoclonal cell lines through scarless genome editing. Genotyping analysis of monoclonal cell lines edited at the *GABPB1L* locus with the indicated pairs of sgRNAs. KO: knockout status (FKO: full knockout; &: heterozygous knockout and/or indels; NA: no detected genotyping bands). *: yellow asterisks highlight clones picked for expansion and further analysis. **A,** HEK293T, *TERT*-promoter wild-type. **B**, HAP1, *TERT*-promoter wild-type. **C**, NHA-S2, *TERT*-promoter wild-type. **D**, LN-18, *TERT*-promoter wild-type. **E**, U-251, G228A *TERT*-promoter mutant. **F**, LN-229, G228A *TERT*-promoter mutant. **G**, T98G, G250A *TERT*-promoter mutant. **H**, Numbers and statistics of monoclonal GABPB1L full knockout cell lines generated. No large indels: clones where no excisions were observed (these clones might still contain small indels at the sgRNA target sites). Het KO: GABPB1L heterozygous knockout (i.e. exon 9 of some but not all *GABPB1L* alleles are excised). FKO: GABPB1L full knockout (i.e. exon 9 of all *GABPB1L* alleles are excised). *: qRT-PCR and western blot analysis revealed that this clone was not a full knockout. Likely, the genome editing removed one or both of the genotyping PCR primer sites.

**Supplementary Fig. 4.**
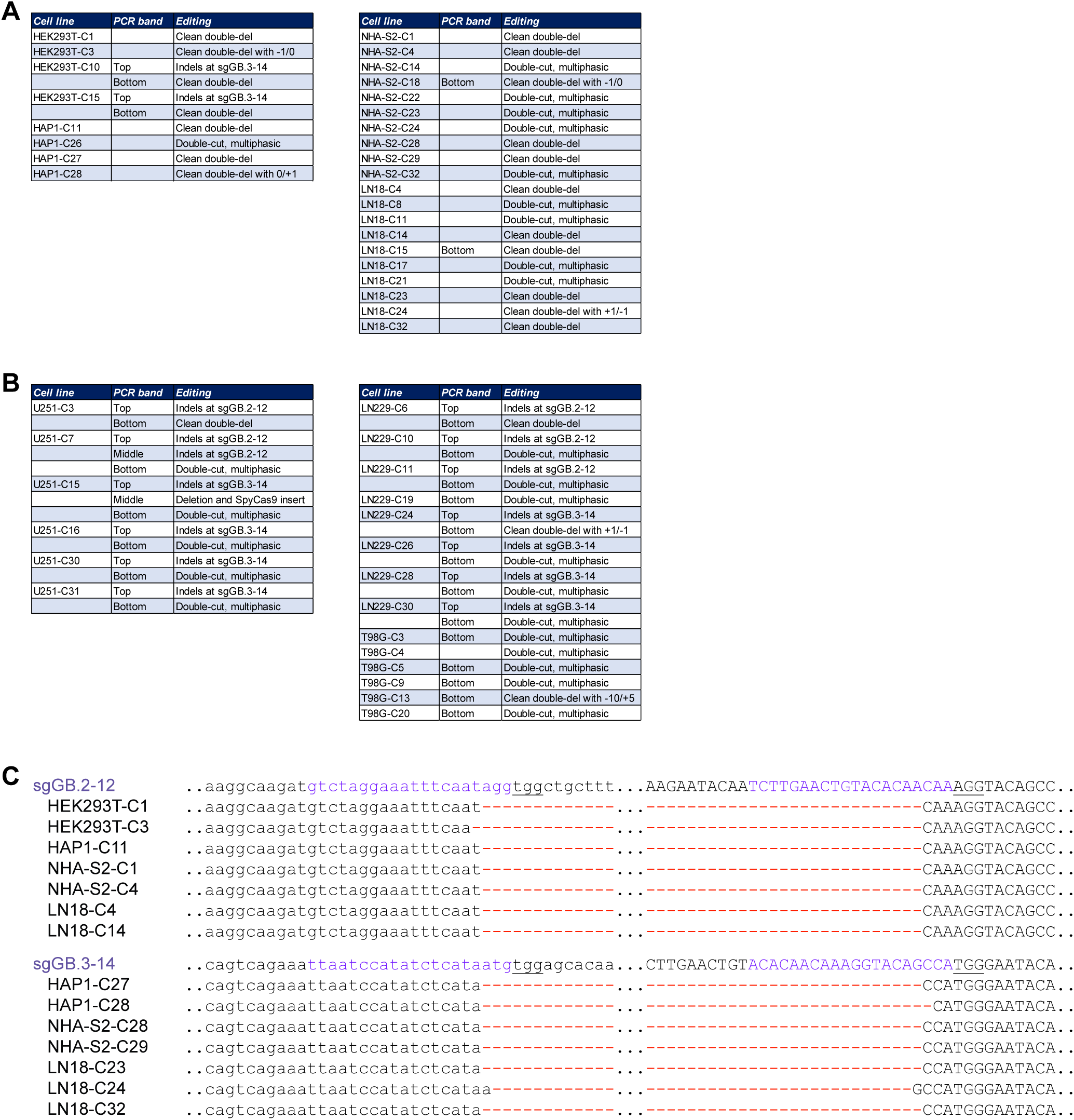
Sequencing-based analysis of monoclonal GABPB1L knockout clones. **A**, Summary of targeted control sequencing of GABPB1L edited monoclonal cell lines. HEK293T, HAP1, NHA-S2 and LN-18 are *TERT*p wild-type cell lines. **B**, U-251, LN-229 and T98G are *TERT*p mutant cell lines. **C**, Genome editing outcome of select full-knockout monoclonal cell lines. sgRNA targeting sites are highlighted in blue. Protospacer adjacent motifs (PAMs) are underlined. Note, most paired-excision cuts are at the expected position between nucleotide -3 and -4 from the PAM.

**Supplementary Fig. 5.**
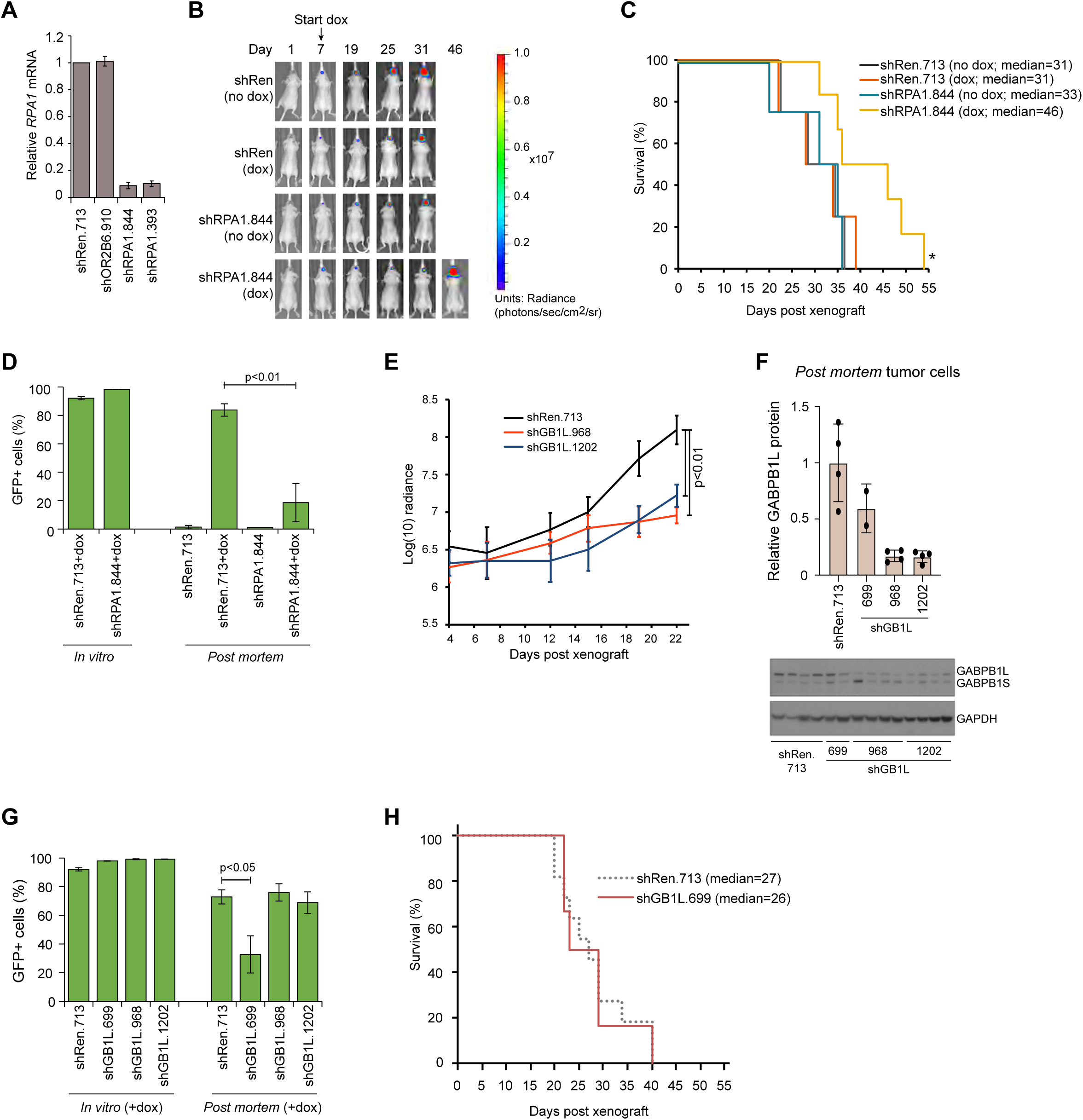
Characterization of inducible shRNAs *in vitro* and *in vivo*. **A**, *RPA1* mRNA expression measured via qRT-PCR in U-251 cells expressing either one of two shRNAs targeting RPA1 (shRPA1.844, shRPA1.393), a negative control (shOR2B6.910), or a non-targeting control (shRen.713). Data are plotted relative to shRen.713. Cells were incubated with doxycycline for 6 days prior to analysis. Data represent mean +/- SEM. **B,C,** Representative bioluminescence images (**B**) and Kaplan-Meier survival curve (**C**) for mice injected with U-251 cells expressing a control shRNA (shRen.713) or RPA1-targeting shRNA (shRPA1.844). Mice were either kept on control chow or fed doxycycline chow to induce shRNA expression (+dox) starting at day 7 post-injection. Doxycycline chow alone had no effect on animal survival. (**C)** *: p-value<0.05 relative to shRen.713 (Kaplan-Meier log-rank test). N = 4-6 mice per condition. **D**, Percentage GFP positive inducible shRNA-expressing U-251 cells measured via flow cytometry analysis as assessed following 3 days of doxycycline incubation in culture (*in vitro*) and following *ex vivo* dissection and dissociation of tumor cells from mice maintained on control chow or doxycycline chow (*post mortem*). **E**, Average log-transformed radiance readings over 23 days of tumor growth for mice bearing orthotopic xenografts of U-251 cells engineered to express doxyxycline-inducible shRNAs targeting either GABPB1L (shGB1L.968, shGB1L.1202) or a non-targeting control (shRen.713). Mice were placed on doxyxycline 7 days post-xenograft. **F**, Immunoblot (bottom) and quantification (top) of GABPB1 protein levels in orthotopic xenograft tumors processed *post mortem* from mice injected with U-251 cells expressing a control shRNA (shRen.713) or GABPB1L-targeting shRNAs (shBG1L.699, shGB1L.968, shGB1L.1202). The lower band represents GABPB1S, the upper band represents GABPB1L **G**, Percentage of GFP positive U-251 cells expressing inducible shRNAs towards GABPB1L or shRen.713 (non-targeting control). GFP was measured via flow cytometry analysis as assessed following three days of doxycycline incubation in culture (*in vitro*) and following *ex vivo* dissection and dissociation of tumor cells from mice maintained on doxycycline chow (*post mortem*). **H**, Kaplan-Meier survival curve displaying median and range of survival for mice injected with U-251 cells expressing an shRNA targeting GABPB1L (shGB1L.699) compared to non-targeting control shRen.713 (dashed line; data also presented in Fig 2G). N = 6 mice per condition for shGABPB1L.699. **D**,**E**,**G**, Data represent mean +/- standard deviation. p: p-value (unpaired, two-tailed student’s t-test).

**Supplementary Fig. 6.**
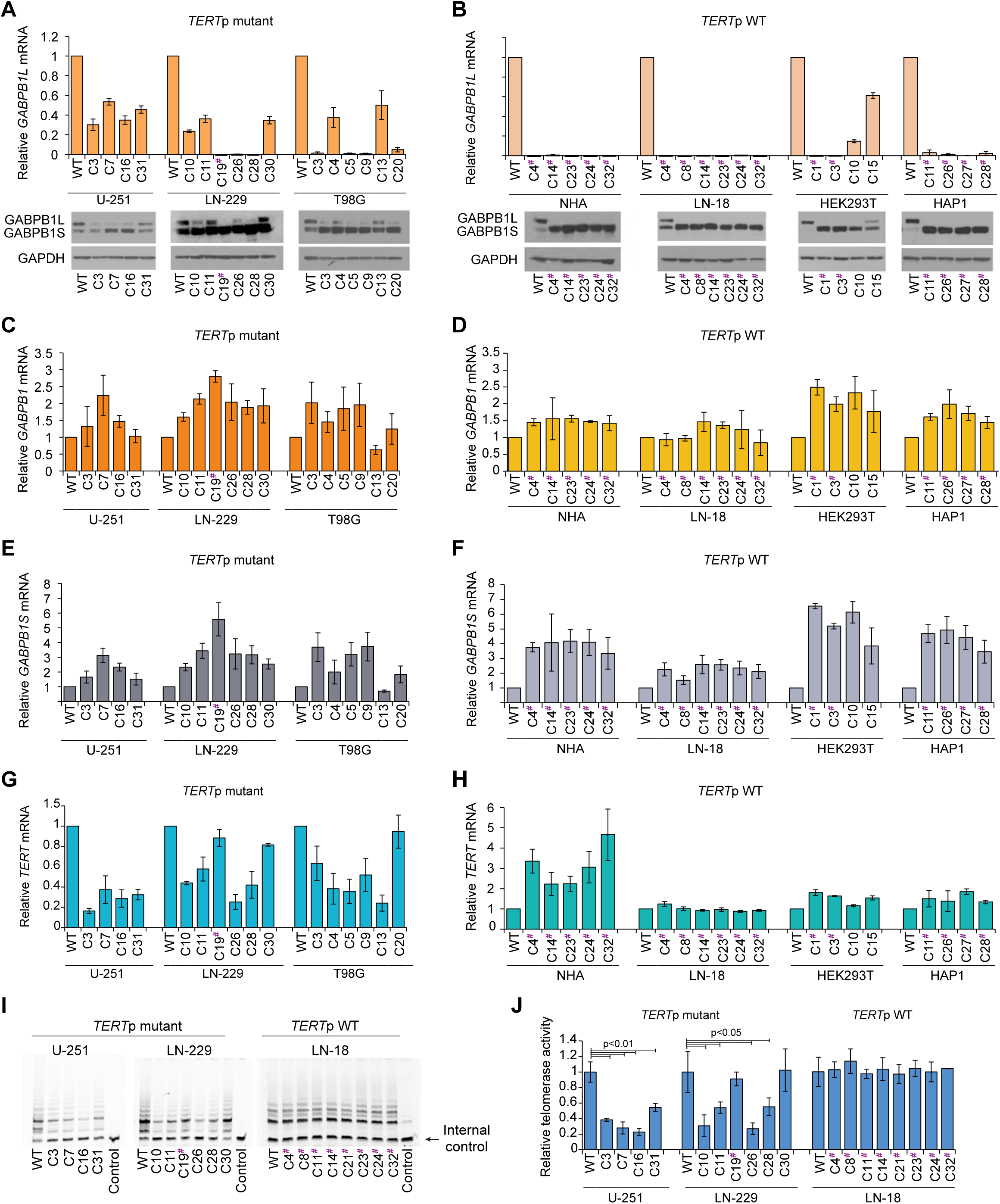
Characterization of GABPB1L knockout cell lines. **A,B**, (Top) Analysis of *GABPB1L* mRNA expression measured via qRT-PCR in GABPB1L knockout clones, plotted relative to control (wild-type) cells. (Bottom) Representative immunoblots of GABPB1 expression in GABPB1L knockout clones compared to control (wild-type) cells. The lower band represents GABPB1S, the upper band represents GABPB1L. Experiments were done in *TERT*p mutant (**A**) and *TERT*p wild-type (**B**) cell lines. **C,D,** Total *GABPB1* mRNA expression measured via qRT-PCR in GABPB1L knockout clones, plotted relative to control (wild-type) cells, in *TERT*p mutant (**C**) and *TERT*p wild-type (**D**) cell lines. **E,F,** *GABPB1S* mRNA expression measured via qRT-PCR in GABPB1L knockout clones, plotted relative to control (wild-type) cells, in *TERT*p mutant (**E**) and *TERT*p wild-type (**F**) cell lines. **G,H,** *TERT* expression measured via qRT-PCR in GABPB1L knockout clones, plotted relative to control (wild-type) cells, in *TERT*p mutant (**G**) and *TERT*p wild-type (**H**) cell lines. **A-G** Data represent mean +/- SEM. **I,J,** Representative gels (**I**) and quantification (**J**) of telomerase activity in GABPB1L knockout clones versus control (wild-type) cells, measured via TRAP assay in 2 *TERT*p mutant cell lines (U-251, LN-229) and 1 *TERT*p WT line (LN-18). Telomerase banding intensity is normalized to internal control band and plotted relative to wild-type. p: p-value (unpaired, two-tailed student’s t-test). Data represent mean +/- standard deviation. **A**-**J**, Hashmarks denote full knockout clones. All others are heterozygous knockout.

**Supplementary Fig. 7.**
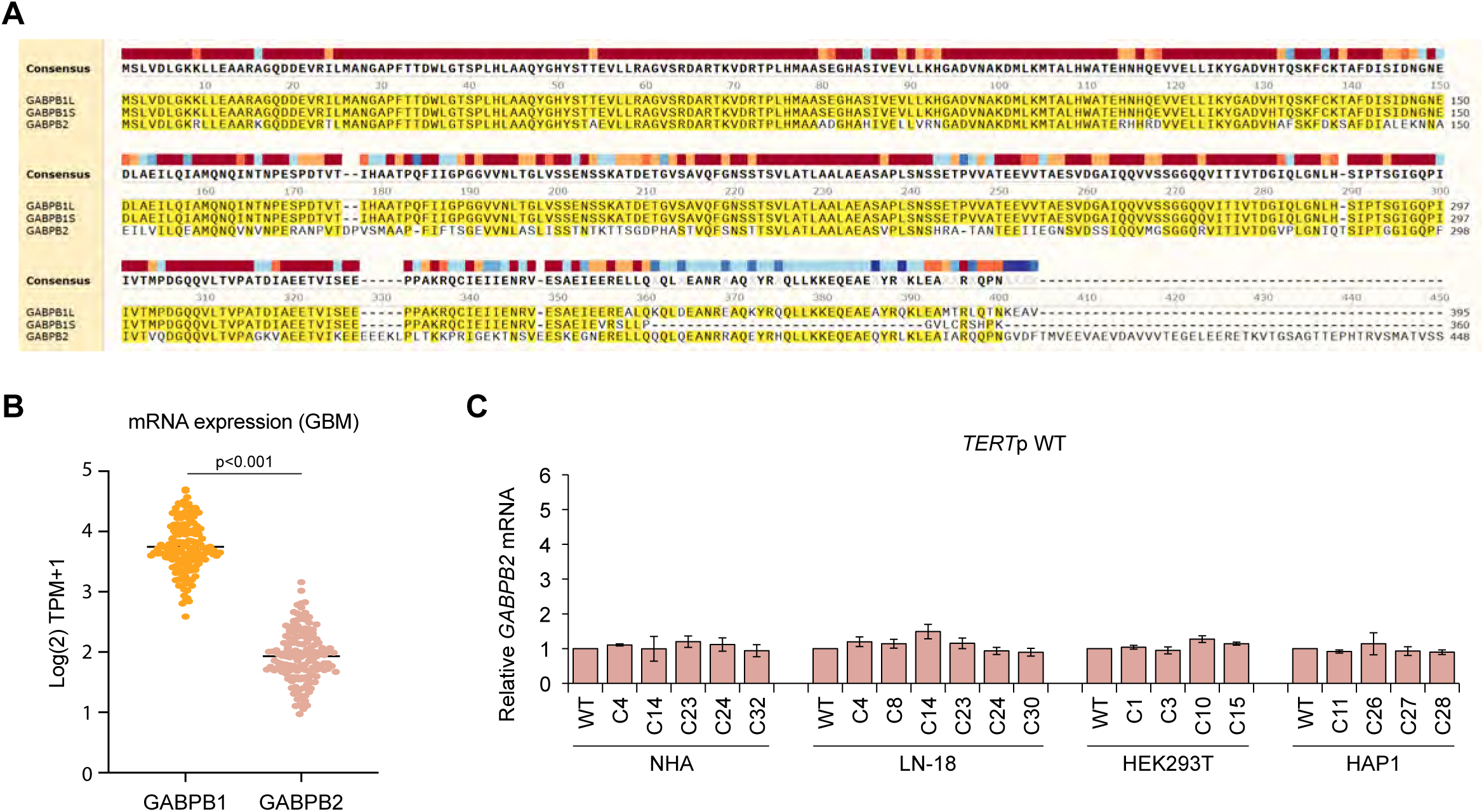
Effect of *GABPB1L* editing on GABPB2. **A**, MUSCLE alignment of GABPB1L, GABPB1S, and GABPB2 protein sequences (SnapGene, version 5.0.7). Matching amino acids are highlighted in yellow. Sequence conservation is shown in colored blocks at the top. **B**, Comparison of GABPB1 and GABPB2 expression levels in GBM patient tumors. Data from the TCGA. p: p-value (Wilcox signed-rank test). **C**, *GABPB2* expression via qRT-PCR in GABPB1L knockout clones plotted relative to control (wild-type) cells in *TERT*p WT cell lines. Data represent mean +/- SEM.

**Supplementary Fig. 8.**
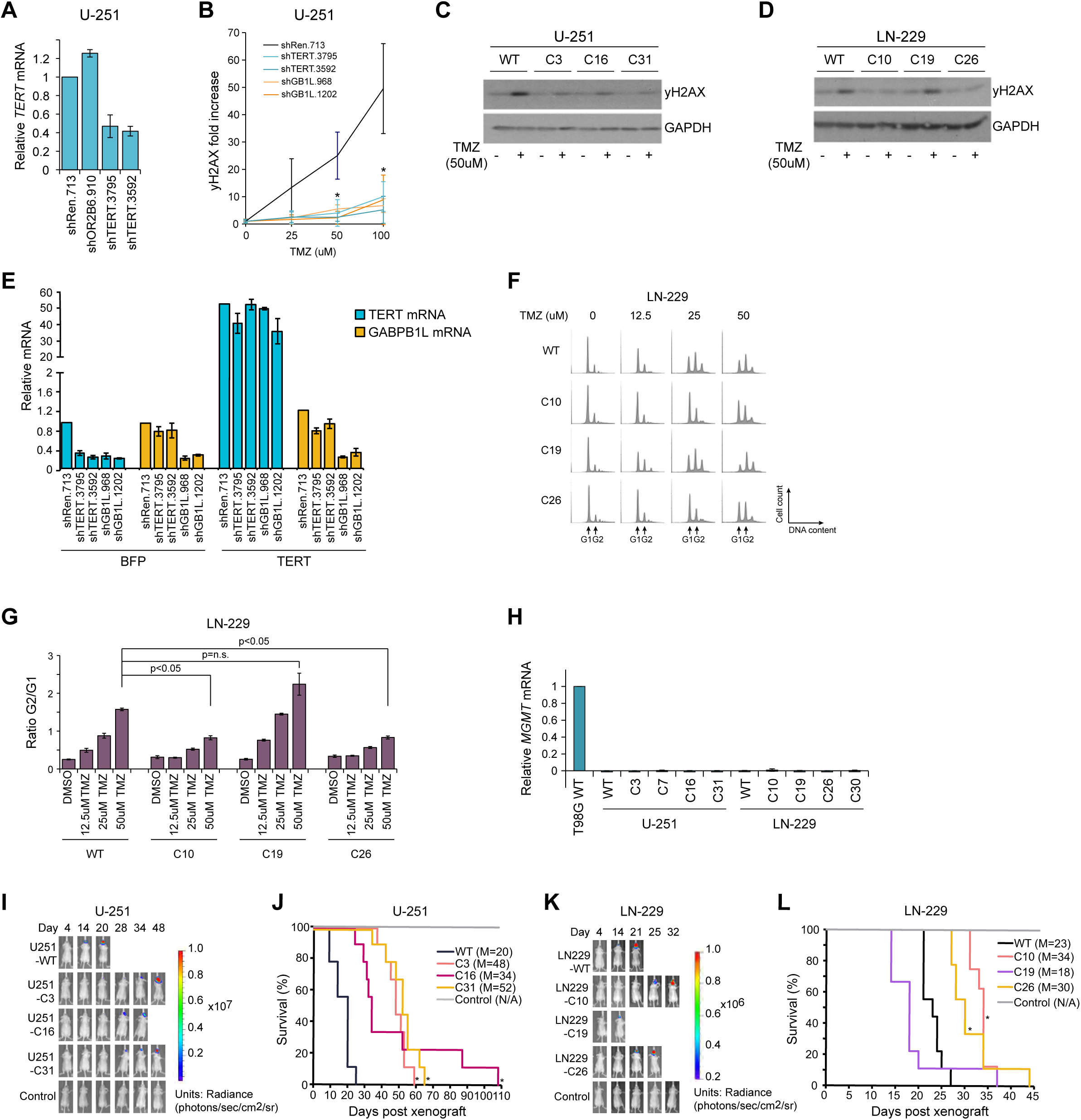
GABPB1L reduction reduces DNA damage response in *TERT* promoter mutant GBM. **A**, *TERT* mRNA expression levels measured via qRT-PCR in U-251 cells with doxycycline-induced shRNAs targeting TERT (shTERT.3795, shTERT.3592) compared to negative control (olfactory receptor OR2B6, shOR2B6.910) and non-targeting (renilla luciferase, shRen.713) shRNAs. Cells were incubated with doxycycline for 6 days prior to harvest. Data represent mean +/- SEM. **B,** Quantification of yH2AX immunoblots in U-251 cells expressing doxycycline-induced shRNAs targeting TERT (shTERT.3795, shTERT.3592) or GABPB1L (shGB1L.968, shGB1L.1202) compared to a non-targeting (renilla luciferase, shRen.713) shRNA. Data normalized to GAPDH and plotted relative to 0 uM TMZ within each condition. *: p-value<0.05 (unpaired, two-tailed student’s t-test). Data represent mean +/- standard deviation. **C**,**D**, Representative immunoblots of wild-type and GABPB1L knockout U-251 (**C**) and LN-229 (**D**) cells treated with 50 uM TMZ for 20 hours prior to harvest. **E**, *TERT* and *GABPB1L* mRNA expression measured via qRT-PCR in U-251 cell lines expressing shRNAs targeting TERT (shTERT.3795, shTERT.3592), GABPB1L (shGB1L.968, shGB1L.1202), or a non-targeting control shRNA (shRen.713), in the prescence of stable expression of BFP (control) or TERT (rescue). Data plotted relative to control (shRen.713, BFP) cells. Data represent mean +/- SEM. **F**,**G**, Representative images (**F**) and G2/G1 ratio quantification (**G**) from FACS-based cell cycle analysis of LN-229 cells treated with a dose titration of TMZ 72 hours prior to harvest. **H**, *MGMT* mRNA expression in U-251 and LN-229 wild-type cells and GABPB1L heterozygous knockout clones, measured via qRT-PCR. MGMT-expressing T98G wild-type cells were used as a positive control. Data plotted relative to T98G wild-type. Data represent mean +/- SEM. **I**,**J**, Representative bioluminescence images (**I**) and Kaplan-Meier survival curves (**J**) for mice injected with wild-type U-251 cells or one of three GABPB1L heterozygous knockout U-251 clones. N = 9 mice per condition. *: p-value<0.01 compared to wild-type (Kaplan-Meier log-rank test). **K,L**, Representative bioluminescence images (**K**) and Kaplan-Meier survival curve (**L**) for mice injected with either wild-type LN-229 cells or one of three GABPB1L knockout LN-229 clones. Note that heterozygous knockout of GABPB1L slowed growth of LN-229 tumors *in vivo*, while the single FKO clone exhibiting GABPB2 upregulation and no change in TERT (LN229-C19) showed unaffected *in vivo* growth. N = 8-9 mice per condition. *: p<0.01 relative to wild-type (Kaplan-Meier log-rank test).

**Supplementary Fig. 9.**
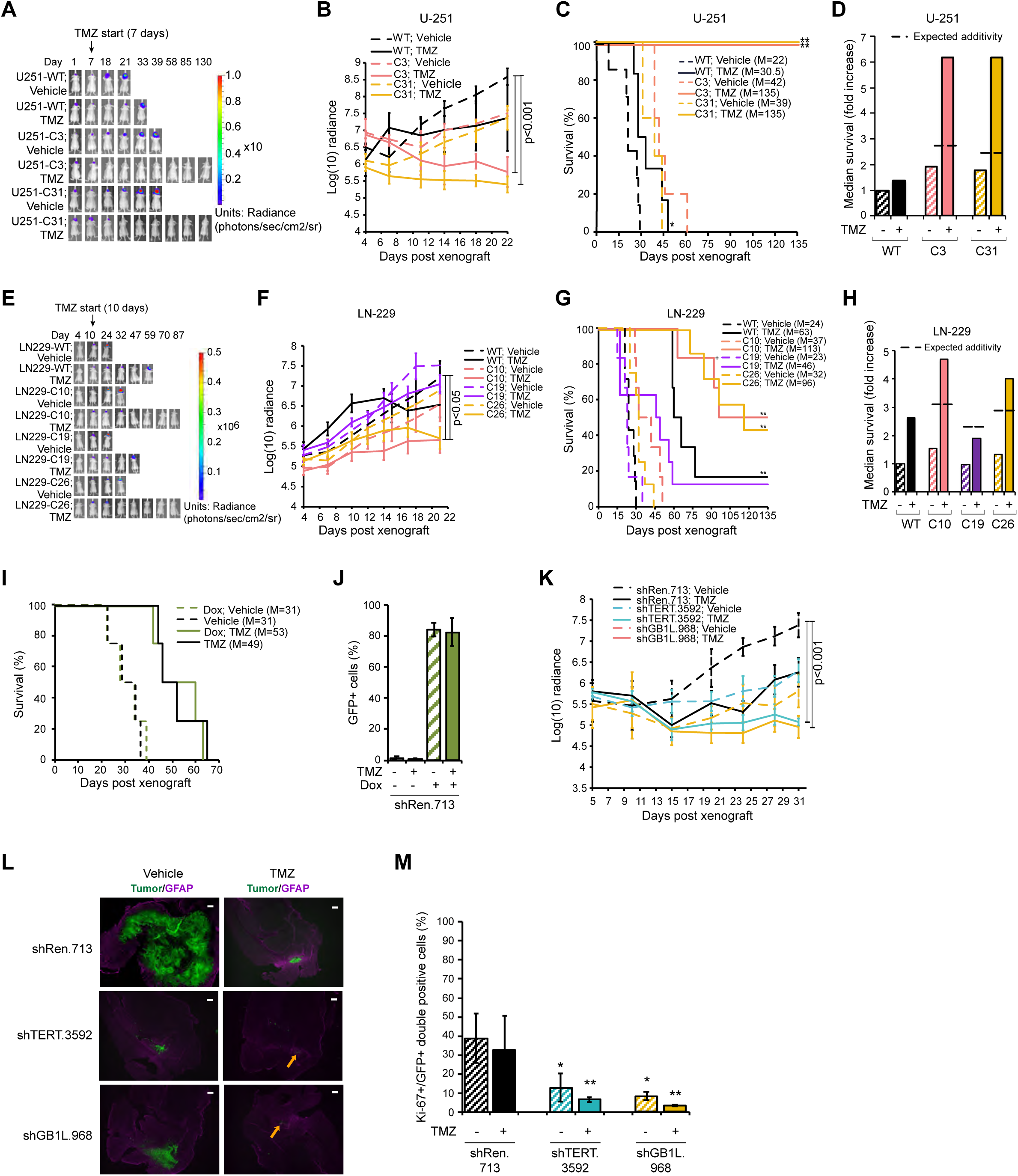
GABPB1L inhibition slows *in vivo* tumor growth and sensitizes tumors to TMZ treatment. **A**,**B**,**C**, Representative bioluminescence images (**A**), average log-transformed radiance readings +/- standard deviation over 22 days of tumor growth (**B**), and Kaplan-Meier survival curves (**C**) for mice injected with either wild-type U-251 cells or one of two GABPB1L heterozygous knockout U-251 clones. Mice were p.o. dosed 7 days post-xenograft with either TMZ or a vehicle control for 10 days (twice 5 days on, 2 days off). N = 6-7 mice per condition. (**B**): p: p-value (unpaired, two-tailed student’s t-test). (**C**): For statistics, survival of mice with no tumor burden was set at the experimental endpoint (135 days). *: p-value<0.05, **: p-value<0.001, relative to wild-type vehicle condition (Kaplan-Meier log-rank test). M: median survival. **D**, Comparison of the relative fold-increase median survival from (**C**) between TMZ alone, GABPB1L heterozygous knockout, and combined TMZ with GABPB1L knockout in U-251 cells. Dashed line represents a simple addition of effects of TMZ and GABPB1L knockout. Survival of mice xenografted with wild-type cells and treated with vehicle control was used for normalization. **E**,**F**,**G**, Representative bioluminescence images (**E**), average log-transformed radiance readings +/- standard deviation over 21 days of tumor growth (**F**), and Kaplan-Meier survival curves (**G**) for mice injected with either wild-type LN-229 cells, one of two GABPB1L heterozygous knockout LN-229 clones (C10, C26), or a full GABPB1L knockout clone (C19). Mice were p.o. dosed with either TMZ or a vehicle control starting 10 days post-xenograft for 5 days. N=6-8 mice per condition. (**F**): p: p-value (unpaired, two-tailed student’s t-test). (**G**): For statistics, survival of mice with no tumor burden was set at the experimental endpoint (135 days). **: p<0.001, relative to wild-type vehicle condition (Kaplan-Meier log-rank test). +: This animal was euthanized due to health problems unrelated to tumor burden. M: median survival. **H**, Comparison of the relative fold-increase median survival between TMZ alone, GABPB1L knockout, and combined TMZ with GABPB1L knockout in LN-229 cells. Dashed line represents a simple addition of effects of TMZ and GABPB1L knockout. Survival of mice xenografted with wild-type cells and treated with vehicle control was used for normalization. **I**, Kaplan-Meier survival curves for mice injected with U-251 cells expressing a doxycycline-inducible control shRNA (shRenilla luciferase), placed on either doxycycline or control chow 7 days post-xenograft, and treated with TMZ or vehicle for 5 consecutive days starting 12 days post-xenograft. Doxycycline administration did not alter the effect of TMZ on survival. M: median survival. **J**, Percentage of GFP positive tumor cells from mice shown in (**I**), measured via flow cytometry analysis, as assessed following *ex vivo* dissociation of tumor cells *post mortem.* Data represent mean +/- standard deviation. **K,** Average log-transformed radiance readings +/- standard deviation over 31 days of tumor growth for mice injected with U-251 cells engineered to express shRNAs targeting TERT (shTERT.3592), GABPB1L (shGB1L.968), or a non-targeting control shRNA (shRen.713). Mice were placed on doxycline chow and p.o. dosed with either TMZ or a vehicle control as described in Figure 4E. **L**, Representative images of immunofluorescence staining in tumors from mice described in Figure 4E, cohort 1. Mice were euthanized 30 days post-orthotopic xenograft for analysis. Doxycycline-induced shRNA-expressing tumor cells are GFP positive. Orange arrows identify the possible tumor region prior to treatment, as identified by scattered GFP-positive cells and signs of glial scarring identified by GFAP staining. N = 3 mice per condition. Scale bar = 300uM. **M**, Quantification of immunofluorescence staining in tumors from mice injected with U-251 cells engineered to express shRNAs towards TERT (shTERT.3592), GABPB1L (shGB1L.968), or a non-targeting control shRNA (shRen.713). Mice were placed on doxycline chow and p.o. dosed with either TMZ or a vehicle control as described in Figure 4E, cohort 1, and euthanized 30 days post-orthotopic xenograft for analysis. N = 5 images per mouse, 3 mice per condition. *: p-value<0.05, **: p-value<0.001 (unpaired, two-tailed student’s t-test).

